# Alternative translation initiation by ribosomal leaky scanning produces multiple isoforms of the Pif1 helicase

**DOI:** 10.1101/2024.01.21.576553

**Authors:** Tomas Lama-Diaz, Miguel G. Blanco

## Abstract

In budding yeast, the integrity of both the nuclear and mitochondrial genomes relies on dual-targeted isoforms of the conserved Pif1 helicase, generated by alternative translation initiation (ATI) of *PIF1* mRNA from two consecutive AUG codons flanking a mitochondrial targeting signal. Here, we demonstrate that ribosomal leaky scanning is the specific ATI mechanism that produces not only these, but also novel, previously uncharacterized Pif1 isoforms. Both in-frame, downstream AUGs as well as near-cognate start codons contribute to the generation of these alternative isoforms. This has crucial implications for the rational design of genuine separation-of-function alleles and provides an explanation for the suboptimal behaviour of the widely employed mitochondrial- (*pif1-m1*) and nuclear-deficient (*pif1-m2*) alleles, with mutations in the first or second AUG codon, respectively. We have taken advantage of this refined model to develop improved versions of these alleles, which will serve as valuable tools to elucidate novel functions of this helicase and to disambiguate previously described genetic interactions of *PIF1* in the context of nuclear and mitochondrial genome stability.

## Introduction

Pif1 belongs to the conserved superfamily 1B (SF1B) of 5’-3’ helicases. In addition to the characteristic SF1B helicase core region, Pif1 and its related proteins also contain a distinctive Pif1 signature motif that has led to the identification of Pif1-like helicases in practically all eukaryotes -including yeasts and human-, eubacteria, archaea and viruses^1^. Although most eukaryotes encode a single Pif1 helicase, *S. cerevisiae* and closely related fungi possess a second paralog, Rrm3, with 40% identical residues to Pif1. In budding yeast, the Pif1 helicase is involved in DNA replication and recombination, safeguarding the integrity of both the mitochondrial and nuclear genomes. The nuclear roles of Pif1 include i) telomere maintenance, as a catalytic inhibitor of telomerase at chromosomal ends and double strand breaks, ii) Okazaki fragment maturation, iii) DNA synthesis during break-induced replication and recombination-dependent replication or iv) support of the replication fork barrier at the rDNA, as reviewed elsewhere^2–4^. To accomplish these functions, Pif1 activities are tightly regulated through protein-protein interactions and post-translational modifications, including lysine acetylation^5^ and checkpoint-dependent phosphorylation^46,7^. Comparatively, its essential function in the maintenance of mitochondrial DNA is poorly understood at the molecular level, but it has been reported that Pif1 travels with the replisome, facilitates repair of DSBs, protects mtDNA from oxidative damage ^8^, and is necessary to maintain iron and zinc homeostasis^9^. To fulfil these roles *PIF1* encodes two different isoforms produced by alternative translation initiation (ATI) from AUG^1^ or AUG^40^, which flank a mitochondrial targeting signal (MTS)^10^. To avoid the pleotropic effects of *PIF1* deletion in genetic studies, two separation-of-function alleles for the mitochondrial (*pif1-m1*) and nuclear (*pif1-m2*) functions were developed, in accordance with the described ATI rationale^10^. In *pif1-m1*, the production of the mitochondrial Pif1 (mPif1) is abolished by mutation of AUG^1^, while nuclear Pif1 (nPif1) can still be translated from the intact AUG^40^ (Supplementary Fig. 1a). Conversely, in *pif1-m2*, the mutation of AUG^40^ abrogates the production of nPif1 without affecting the translation of mPif1 (Supplementary Fig. 1b). *pif1-m2* has been widely exploited to interrogate the involvement of Pif1 in nuclear DNA replication and repair. However, since its first description, *pif1-m2* mutants have been acknowledged to retain a certain amount of nuclear protein, as inferred from their partial suppression of some of the *pif1Δ* phenotypes in different genetic set-ups^11–17^.

Here we demonstrate that ribosomal leaky scanning is the main ATI mechanism underlying start codon selection in *PIF1* mRNA and characterize a novel alternative nuclear isoform of Pif1. Moreover, we uncover the ability of near-cognate AUG codons to produce a phenotypically relevant amount of functional Pif1. These previously unrecognized sources of Pif1 activity may explain not only the residual nuclear activity of *pif1-m2*, but also the less explored incomplete phenotype of *pif1-m1* mutants. This refined model, together with the identification of Pif1 nuclear localization signal, has allowed us to develop improved nuclear- and mitochondrial-deficient alleles for Pif1, which can be employed to disambiguate misleading genetic interactions.

## Results

### The *pif1-m2* allele does not fully recapitulate the phenotypes due to loss of nuclear Pif1 functions in *pif1Δ* mutants

The study of Pif1 roles in both nuclear and mitochondrial genome stability has taken advantage of the two classical separation-of-function mutants *pif1-m1* and *pif1-m2*^10^. However, results from several groups^11–17^ and our own suggest that *pif1-m2* retains variable levels of nuclear activity. To assess the relevance of this residual nuclear activity, we compared directly the phenotypes derived from *pif1-m2* and *pif1Δ* for some of the described genetic interactions. Importantly, in our experiments we also included the *pif1-m1* mutant to clarify whether any phenotypic differences between *pif1-m2* and *pif1Δ* arise from the residual nuclear activity in *pif1-m2* or the combined effect of Pif1 loss at both nucleus and mitochondria (Fig.1). First, we tested the synthetic sickness between *pif1Δ* and the absence of *DIA2*, a member of the E3 ligase SCF complex^18^ by tetrad dissection analyses of *DIA2/dia2Δ* heterozygous diploids in combination with *PIF1*, *pif1Δ*, *pif1-m1* or *pif1-m2* (Fig.1a). Neither *pif1-m1* or *pif1-m2* conferred an overt synthetic sickness phenotype, as observed in *pif1Δ dia2Δ* double mutants, and consistently with a previous report^19^. Similarly, the hypersensitivity to HU in *pif1Δ rad3-102* (a hypomorphic allele of the essential Rad3 helicase involved in nucleotide excision repair) double mutants^20^, could not be recapitulated with the *pif1-m1* or *pif1-m2* mutants (Fig. 1b). We next compared how these separation-of-function alleles could suppress the lethality caused by the absence of the essential Dna2 nuclease-helicase (Fig. 1c). Consistent with the literature^21^, the complete absence of Pif1 provides a robust suppression of *dna2Δ* lethality that is not observed with the *pif1-m1* allele. However, *pif1-m2 dna2Δ* cells grow comparatively poorly to the *pif1Δ dna2Δ* colonies, indicative of reduced, but functionally relevant amounts of nuclear Pif1 in those cells. In this sense, replacement of the endogenous *pif1-m2* promoter with the constitutive *ADH1* promoter is sufficient to restore *dna2Δ* lethality (Fig. 1d), suggesting that functional nuclear Pif1 can still be produced from the *pif1-m2* mRNA. Interestingly, the synthetic lethality observed between *pif1Δ* and *top3Δ*^22^ is fully recapitulated by *pif1-m2* (but not *pif1-m1*), suggesting that nuclear levels of Pif1 in *pif1-m2* cells are too low to prevent lethality (Fig. 1e). In this case, the replacement of the endogenous promoter of *pif1-m2* with *P_ADH1_* is sufficient to suppress this synthetic lethality (Fig. 1f), in accordance with increased Pif1 levels being able to ameliorate the slow growth of *top3Δ* mutants^22^. Finally, given Pif1 role in preventing aberrant telomeric expansions^10,23^, we analysed telomeric length in these mutants. In agreement with the presence of functional nuclear Pif1 in *pif1-m2* mutants, we recapitulated the previously observed^23^ intermediate telomere length of these mutants compared to wild-type and *pif1Δ* strains (Fig. 1g).

**Fig 1.**
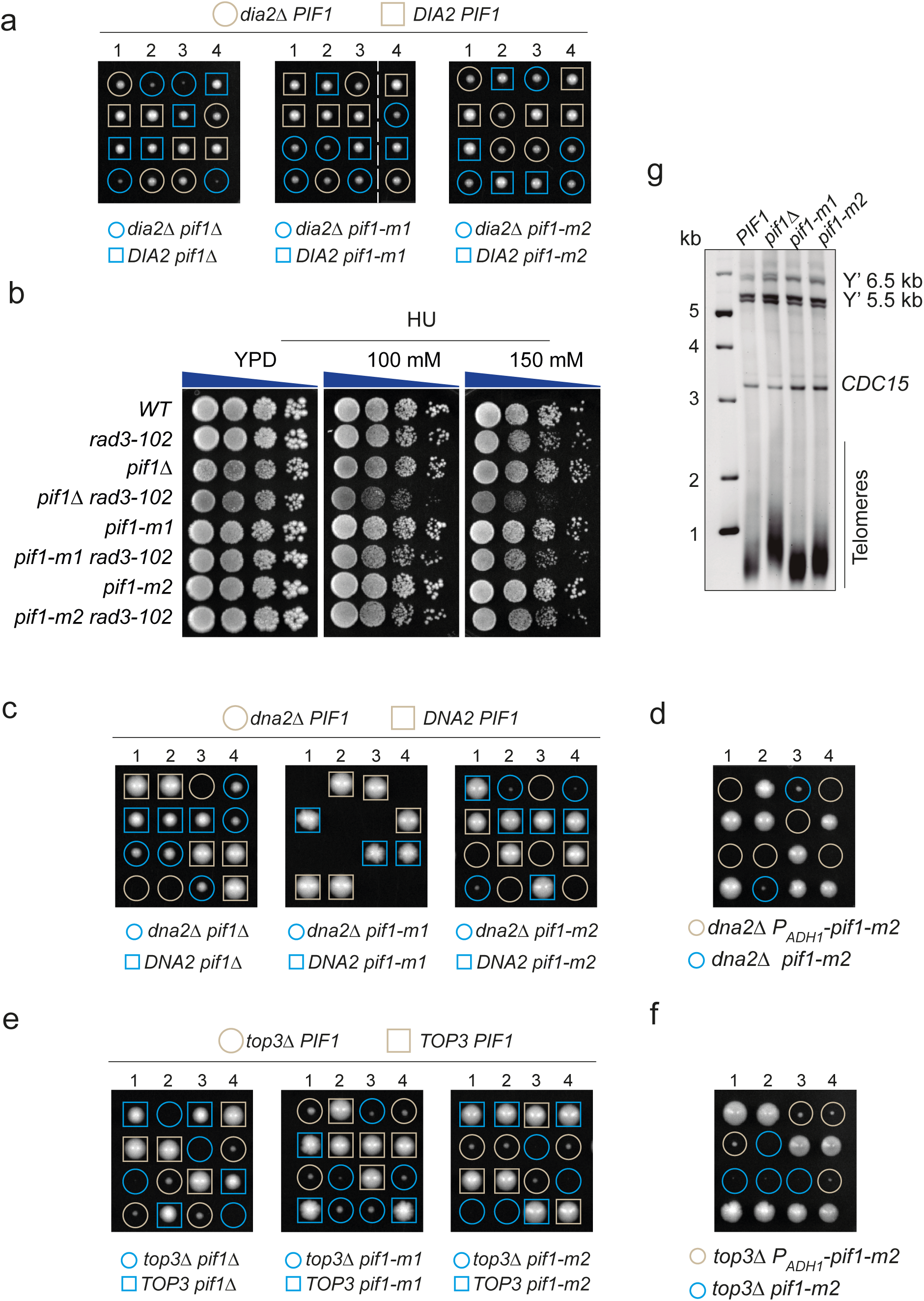
*pif1-m1* and *pif1-m2* cannot disambiguate if the mitochondrial or nuclear loss of Pif1 function underlies various *pif1Δ* genetic interactions. (a) Neither *pif1-m1* or *pif1-m2* recapitulate the synthetic sickness between *pif1Δ* and *dia2Δ*. Tetrad microdissection of diploid strains carrying heterozygous mutations for the indicated wild-type and mutant alleles of *DIA2* and *PIF1*. Images were taken after 2 days of incubation at 30 °C. Dashed lines indicate the stitching of two images from the same plate. (b) Neither *pif1-m1* or *pif1-m2* recapitulate the hypersensitivity to HU of *pif1Δ* and *rad3-102*. Ten-fold serial dilutions of strains with the indicated genotypes were plated on YPD containing different concentrations of hydroxyurea (HU) and imaged after 2 days of incubation at 30 °C. (c) *pif1-m2* behaves as a hypomorphic allele for the loss of Pif1 nuclear function in the context of the suppression of *dna2Δ* lethality. As in (a), but employing heterozygous strains for the indicated wild-type and mutant alleles of *DNA2* and *PIF1*. Images were taken after 3 days of incubation at 30 °C. (d) Overexpression of *pif1-m2* is lethal in *dna2Δ* cells. As in (c), but employing the constitutively overexpressed allele *P_ADH1_ -pif1-m2*. (e) *pif1-m2* recapitulates the synthetic lethality between *pif1Δ* and *top3Δ*. As in (c), but employing heterozygous strains for the indicated wild-type and mutant alleles of *TOP3* and *PIF1*. (f) Overexpression of *pif1-m2* sustains growth of *top3Δ* cells. As in (e), but employing the constitutively overexpressed allele *P_ADH1_ -pif1-m2*. (g) Neither *pif1-m1* or *pif1-m2* drive telomere extension to the extent of *pif1Δ*. Southern blot of XhoI-digested DNA from strains carrying the indicated *PIF1* alleles using the subtelomeric Ý probe (see *Methods*). Telomeric DNA can be observed as a smeared band at the bottom of the blot. A second probe targeted to *CDC15* was employed as a loading control.

Altogether, these results demonstrate that the separation-of-function allele *pif1-m2* still retains a certain level of nuclear activity that may be sufficient to mask some of the phenotypes observed in the complete absence of *PIF1*. Moreover, this also implies that in those *pif1Δ* interactions not recapitulated by *pif1-m1* or *pif1-m2*, it cannot be formally disambiguated whether this is a consequence of the incomplete separation-of-function of these mutant alleles or whether such interactions require the simultaneous loss of nuclear and mitochondrial Pif1 functions.

### An additional nuclear isoform of Pif1 produced by ATI is increased in *pif1-m2* mutants

The mitochondrial and nuclear isoforms of Pif1 are produced through the alternative translation initiation of *PIF1* mRNA at two different, in-frame AUG codons, corresponding to methionines at positions 1 and 40 of Pif1 (Fig. 2a). The full-length isoform contains an N-terminal mitochondrial targeting signal (MTS) that drives its import into mitochondria^10^. Once in the mitochondrial matrix, the full-length protein is proteolytically processed to yield the mature mitochondrial Pif1 (mPif1) starting at arginine 46^24^. Alternatively, when translation starts at AUG^40^, the MTS is excluded and the protein is imported into the nucleus (nPif1), presumably through a yet-uncharacterized nuclear localization signal (Fig. 2a). This renders the mature nPif1 (∼93 kDa) just fractionally bigger than the mature mPif1 (∼92 kDa) and, therefore, their electrophoretic mobility is very similar. In *PIF1* cells both isoforms form a tight doublet when analysed by western blotting, although their different migrations are more apparent in the *pif1-m1* and *pif1-m2* mutants (Fig. 2b). We noticed the presence of a mild, but specific, third band in all *PIF1* alleles and in two different genetic backgrounds, displaying faster electrophoretic mobility and being particularly enriched in *pif1-m2* mutants (Fig. 2b, red arrow). In these cells, this band is over 3 times more abundant than in *PIF1* or *pif1-m1* cells, representing around 30% of total Pif1 (Fig. 2c and Supplementary Fig. 2a).

**Fig 2.**
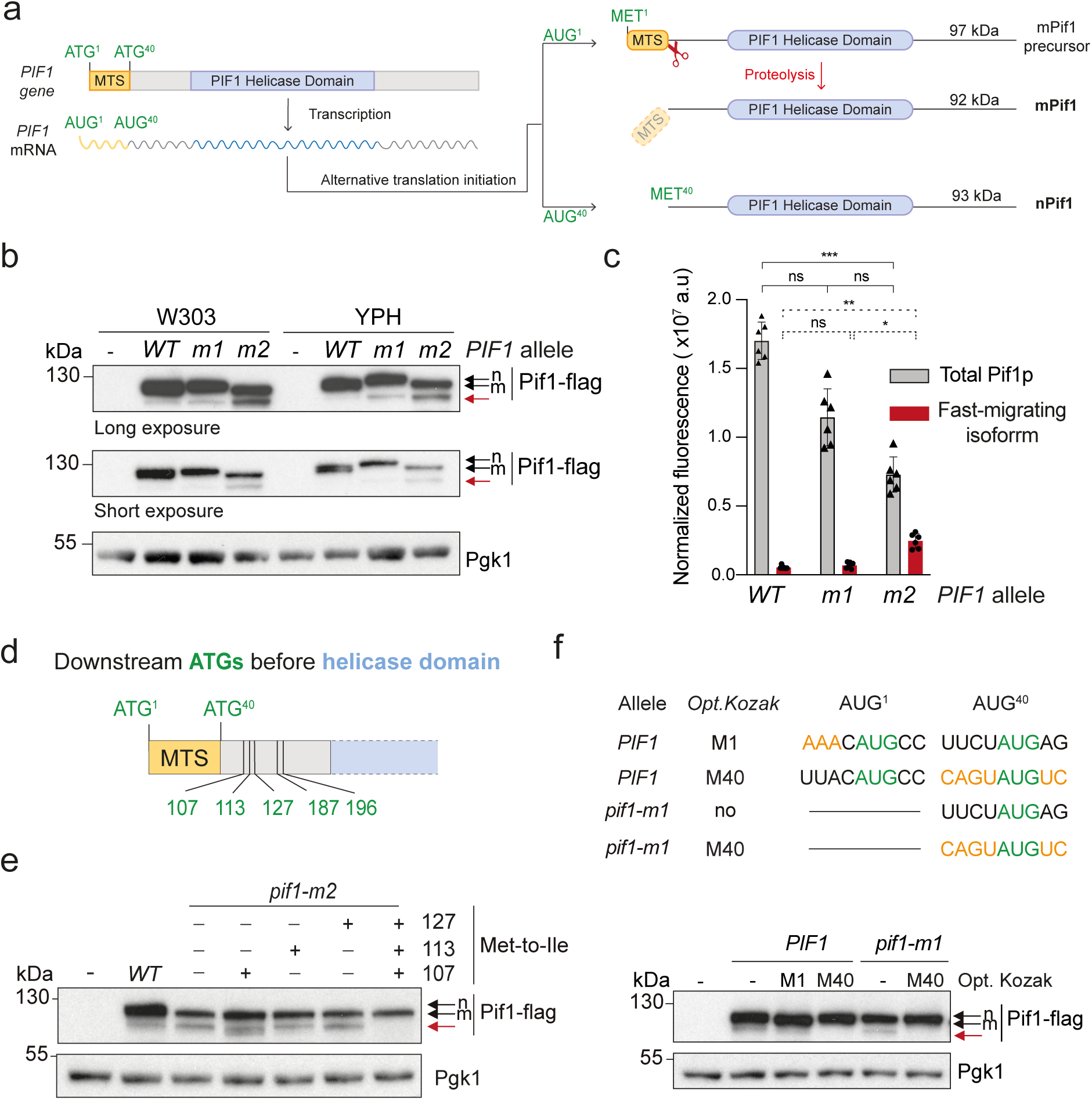
Alternative translation initiation of *PIF1* mRNA by ribosomal leaky scanning drives the production of a novel Pif1 isoform. (a) Schematic representation of the current model for Pif1 isoforms productions by ATI. (b) Detection of a novel, fast-migrating Pif1 isoform. Protein extracts from strains expressing untagged *PIF1* (-), *PIF1-6flag* (*WT*), *pif1-m1-6flag* (*m1*) and *pif1-m2-6flag* (*m2*) strains were probed with an anti-FLAG antibody to assess Pif1 expression (two different exposures shown). Black arrows indicate the tight doublet formed by nuclear (n) and mitochondrial Pif1 isoforms (m). Red arrows mark the fast-migrating isoform. Pgk1 was employed as a loading control. (c) The fast-migrating isoform of Pif1 is significantly enriched in *pif1-m2* mutants. Quantification of total Pif1 or the fast-migrating isoform in *PIF1-6flag*, *pif1-m1-6flag* and *pif1-m2-6flag* strains from fluorescent western blots (see Suppl. Fig. 2). Histograms represent mean values ± standard deviation (n = 6). Statistical analysis was performed using ANOVA Kruskal-Wallis test followed by Dunn’s post hoc multiple comparison analysis. P-value represented by ns, *p<0.05, ** p<0.005, and *** p<0.005. (d) Schematic representation of the 5’ region of *PIF1*, indicating the in-frame ATG codons before the helicase domain. (e) Simultaneous mutation of Met codons to Ile at positions 107, 113 and 127 abrogates the production of the fast-migrating isoform. Western blot analysis of Pif1 in wild-type of *pif1-m2* strains harbouring the indicated single and triple Met to Ile substitutions. Arrows as in (b). (f) Optimization of the Kozak context surrounding AUG^40^ in *PIF1* and *pif1-m1* abrogates the production of Pif1^107–859^. Western blot analysis of Pif1 was carried out as in (e). The specific mutations introduced to optimize the Kozak context of start codons (green) AUG^1^ and AUG^40^ in each *PIF1* allele are shown in orange.

We speculated that this band could arise from an additional alternative translation initiation event downstream AUG^40^. There are five in-frame AUG codons between this position and the beginning of the Pif1 helicase domain, clustered in positions corresponding to the methionines 107, 113 and 127, and methionines 187 and 196 (Fig. 2d). Initiation of translation from any of these positions would result in a Pif1 isoform that would retain an intact helicase domain but would lack the N-terminal region between M40 and this alternative start site. To test this hypothesis, we ectopically expressed *PIF1* alleles (Supplementary Fig. 2b) with single conservative methionine-to-isoleucine mutations at each of these positions and determined whether the expression of the fast-migrating isoform was affected, but no obvious changes were observed (Supplementary Fig. 2c). However, since an ectopically expressed Pif1^107–889^ truncation co-migrates electrophoretically with the fast-migrating isoform (Supplementary Fig. 2d), we decided to focus on the M107-M113-M127 cluster for further analysis (Fig. 2d). Interestingly, double Met-to-Ile substitutions in this cluster revealed that translation initiation may occur from any of these methionines, albeit M107 appears as the most prominent one (Supplementary Fig. 2e). In this sense, a fainter band with further increased mobility can be observed when only M127 is maintained (Supplementary Fig. 2e). Importantly, the fast-migrating band became undetectable when these three methionines were simultaneously mutated (Fig. 2e). These results demonstrate the existence of, at least, a third cellular Pif1 isoform that is translated from M107 (hereinafter, Pif1^107–859^) and that M113 and M127 can also function as alternative initiation points.

### The nuclear Pif1 isoforms are produced through ribosomal leaky scanning

Several mechanisms of alternative translation initiation have been proposed to explain the generation of different protein isoforms from a single mRNA, including internal ribosomal entry sites, reinitiation, ribosomal shunting and ribosomal leaky scanning^25,26^. Leaky scanning involves the ribosomal skipping of the first AUG codon and initiation of translation at a downstream, in-frame AUG codon^27^. In this context, the efficiency of any given AUG for translation initiation depends on the sequence context around this codon, known as the Kozak sequence. Interestingly, the Kozak sequence for AUG^1^ in *PIF1* mRNA is a relatively poor context for initiation (Suppl. Fig. 2f), consistent with a model in which ribosomes must have a high probability of skipping AUG^1^ (that produces mPif1) in order to reach AUG^40^ and produce nPif1. Since the Kozak context around AUG^40^ is also sub-optimal, we could further extend this model and hypothesize that a fraction of ribosomes skipping AUG^40^ reach AUG^107^ and promote translation of Pif1^107–859^. If true, we would predict that reducing the leakiness of ribosomal scanning over AUG^40^ should decrease the translation of Pif1^107–859^. Therefore, we designed optimized, but conservative, Kozak contexts around AUG^1^ or AUG^40^ in the *PIF1* and *pif1-m1* alleles and verified the expression of Pif1^107–859^ by western blotting (Fig. 2f). Confirming our hypothesis, no Pif1^107–859^ could be detected in those cells harbouring constructs with optimized Kozak contexts at AUG^40^. Also, optimization of AUG^1^ in *PIF1* cells produced an apparent increase in mobility of the Pif1 doublet that would be consistent with increased proportion of mPif1 over nPif1. Altogether, our data demonstrate that the mechanism underlying the alternative translation of Pif1 isoforms is ribosomal leaky scanning and explain the specific increase of Pif1^107–859^ in *pif1-m2* mutants, since the mutation of AUG^40^ in this allele would allow a higher proportion of scanning ribosomes to reach AUG^107^.

### Pif1^107–859^ is catalytically active and retains partial functionality in the nucleus

Pif1^107–859^ lacks the MTS region and, consequently, cells expressing this allele display a *petite* phenotype and are unable to grow on non-fermentable carbon sources (Supplementary Fig. 3a). Hence, to investigate if the cause of the incomplete separation-of-function of *pif1-m2* alleles is due to the increased levels of nuclear Pif1^107–859^, we first decided to assess its subcellular localisation by fluorescence microscopy. For this purpose, we integrated constructs carrying *PIF1-eGFP* alleles under the control of the inducible *GAL1* promoter in a strain with endogenous, untagged *pif1-m2* (to avoid the confounding effect of the *petite* phenotype). Upon induction, Pif1-eGFP signal could be observed both in the nucleus and mitochondria in cells expressing *PIF1*, while in those expressing *pif1-m1* or *pif1^107–859^* the Pif1 staining was highly enriched in the nucleus (Fig. 3a). Cells expressing *pif1-m2-eGFP*, in addition to the predicted mitochondrial localization, also displayed a mild Pif1 nuclear enrichment (Fig. 3a), in accordance with the genetic data in this paper and the literature. We next determined whether Pif1^107–859^ retained catalytic activity with respect to the wild-type protein. For this purpose, we expressed Pif1, Pif1^107–859^ and the catalytically inactive Pif1^KA^ ^23^ as C-terminal His-tagged fusion proteins in *E.coli* (Supplementary Fig. 3b) and compared their ATPase (Supplementary Fig. 3c and d) and helicase (Fig. 3b and Supplementary Fig. 3e) activities. Our results indicate that the truncation of the N-terminal region does not impair either Pif1^107–859^ ATPase or helicase activity with respect to wild-type Pif1.

**Fig 3.**
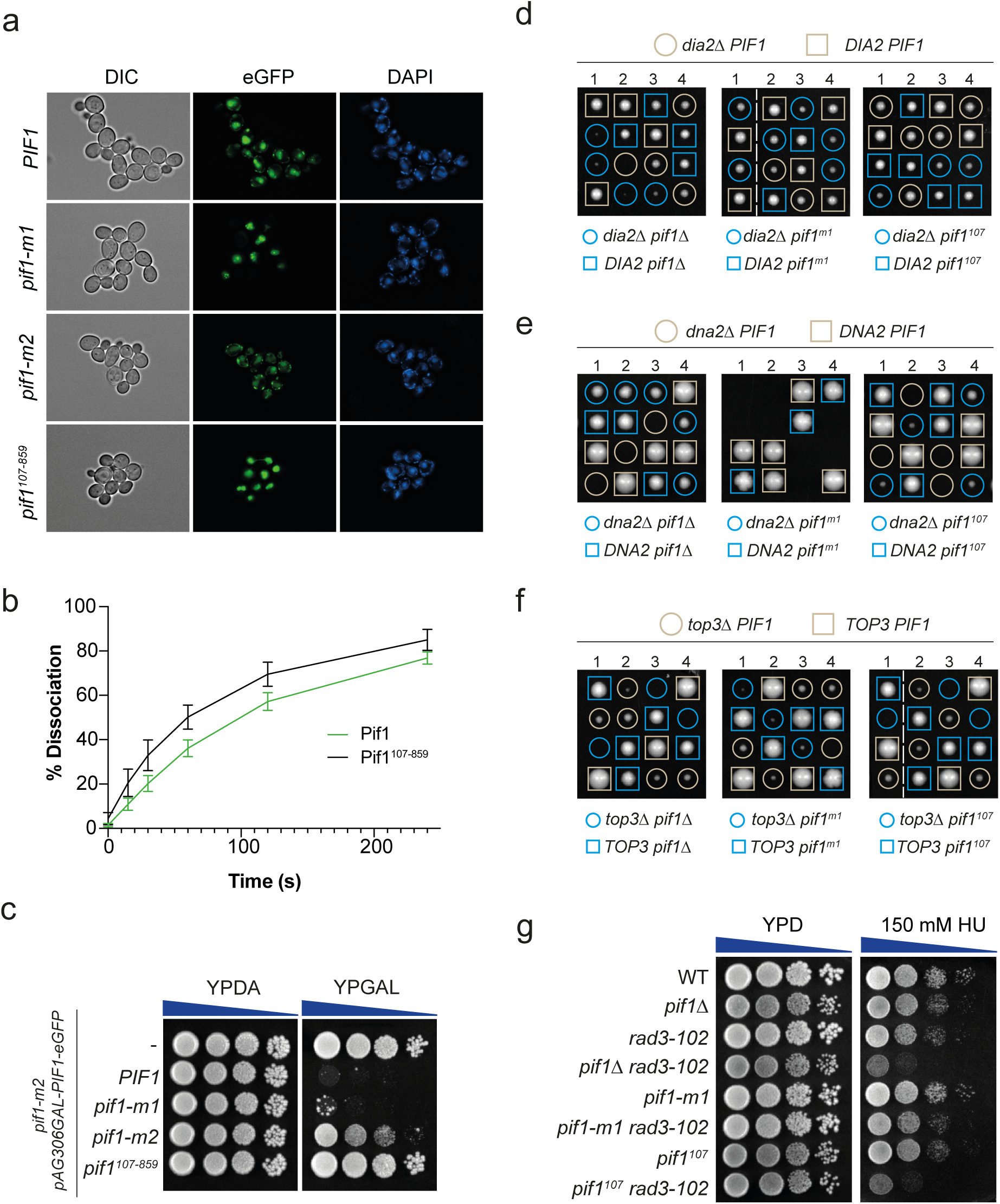
Pif1^107–859^ is a source of catalytically active Pif1 in the nucleus. (a) Pif1^107–859^ is a nuclear Pif1 isoform. Subcellular localization of Pif1 isoforms in strains overexpressing different *PIF1-eGFP* alleles was assessed in live cells by fluorescence microscopy. DNA was visualized with DAPI staining and cell contour by DIC microscopy. (b) Helicase activity of Pif1^107–859^ *in vitro* is comparable to wild-type Pif1. DNA unwinding activity of purified Pif1 and Pif1^107–859^ was estimated as percentage of substrate dissociation in time-course assays from three replicates (see Supp. Fig. 3 for representative image) and represented as mean values ± standard deviation. (c) Overexpression of *pif1*^107–859^ is not lethal. Ten-fold dilutions of strains with the indicated genotypes were plated on YPD or YPGal plates and imaged after 2 days of incubation at 30 °C. (d) *pif1^107–859^*suppresses synthetic sickness of *dia2Δ pif1Δ* double mutants. Tetrads from diploid strains carrying heterozygous mutations for the indicated wild-type and mutant alleles of *PIF1* and *DIA2* were microdissected on YPD plates and incubated at 30 °C for 2 days before imaging. (e) *pif1^107–859^*is not as efficient as *pif1Δ* in suppressing the lethality of *dna2Δ*. Tetrads from diploid strains carrying heterozygous mutations for the indicated wild-type and mutant alleles of *PIF1* and *DNA2* were microdissected on YPD plates and incubated at 30 °C for 3 days before imaging. (f) *pif1^107–859^*is synthetic lethal with *top3Δ*. Tetrads from diploid strains carrying heterozygous mutations for the indicated wild-type and mutant alleles of *PIF1* and *TOP3* were microdissected on YPD plates and incubated at 30 °C for 3 days before imaging. (g) *pif1^107–859^*recapitulates the hypersensitivity to HU of *pif1Δ* in *rad3-102* mutants. Ten-fold serial dilutions of strains with the indicated genotypes were plated on YPD in the presence or absence of hydroxyurea (HU) and imaged after 2 days of incubation at 30 °C.

To understand if this nuclear and catalytically proficient isoform could substitute for full-length nPif1, we first addressed the biological functionality of Pif1^107–859^ using as a proxy the lethality caused by the overexpression of nuclear Pif1^5,28,29^. While the overexpression of *PIF1* or *pif1-m1* derived in extensive cell death, overexpression of *pif1*^107–859^ caused no detectable reduction in cell viability (Figure 3c), despite their comparable protein levels (Supplementary Fig. 3f), suggesting that the N-terminal region in Pif1 may be relevant for some Pif1 functions *in vivo*.

Given these results, we directly addressed whether Pif1^107–859^ could be responsible for incomplete loss of Pif1 nuclear function in *pif1-m2* mutants. First, tetrad dissection analyses showed that *dia2Δ pif1^107–859^* mutants displayed reduced synthetic sickness compared to *dia2Δ pif1Δ* mutants (Fig. 3d), indicating that Pif1^107–859^ is able to support cell growth similarly to Pif1 in *dia2Δ* mutants. Second, unlike the lethality caused by the absence of *DNA2* in *PIF1* and *pif1-m1* strains, *dna2Δ pif1^107–859^* mutants are able to grow, but form smaller colonies than *dna2Δ pif1Δ* mutants, indicating that in this context Pif1^107–859^ is partially functional (Fig. 3e). Third, *top3Δ pif1^107–859^*strains failed to grow after tetrad dissection, similarly to *top3Δ pif1Δ* strains, suggesting that Pif1^107–859^ cannot substitute for Pif1 in these conditions (Fig. 3f). Fourth, *pif1^107–859^* could not reduce the hypersensitivity to HU observed in *rad3-102 pif1Δ* mutants, unlike *pif1-m1* (Fig. 3g), an unexpected result that could arise due to various reasons and will be further discussed below. Taken together, our data indicate that in *pif1-m2* mutants there is an increase in the translation of the Pif1^107–859^ isoform, which is able to enter the nucleus and fulfil some of the functions of nuclear Pif1.

### Identification of the nuclear localization signal in Pif1

The presence of alternative nuclear Pif1 isoforms may lead to the misinterpretation of genetic results obtained with the classical *pif1-m2* allele. Therefore, we argued that a more robust separation-of-function allele could be attained by blocking the nuclear import of all isoforms. While it has been proposed that Pif1 contains a nuclear localization signal (NLS)^30^, this had not been formally shown at the time this project was started. Therefore, we employed an *in silico* analysis of the Pif1 sequence to search for potential NLSs and selected four candidate regions for further analysis (Supplementary Fig. 4a, see also *Methods*). As a first approach to test their functionality, we took advantage of the *P_GAL1_-PIF1* overexpression-dependent lethality to search for those regions whose deletion restored viability upon *GAL1* promoter induction (Supplementary Fig. 4b). We observed that deletion of two different regions (650-659 and 775-803) resulted in an increased survival, suggesting that both regions might be involved in Pif1 nuclear import. However, fluorescence microscopy of C-terminal eGFP-fusions of these *PIF1* variants revealed that only deletion of the 775-803 region resulted in loss of nuclear accumulation of Pif1, concomitantly with an increase in cytoplasmic signal (Supplementary Fig. 4c). These results indicate that the 775-803 region is required for the nuclear import of Pif1, while the 650-659 deletion probably results in a general loss-of function of the protein. To pinpoint the key residues in this NLS, we made substitutions to alanine in two patches of basic residues present in this region, (Fig. 4A) and determined whether these mutations affected functionality of this NLS by fluorescence microscopy (Fig. 4b) and suppression of overexpression-dependent lethality of *P_GAL1_-PIF1* (Fig. 4c-d). Substitution of all nine basic aminoacids (Pif1^9A^) in this region or exclusively in the first basic patch (^781^KKRK^784^; Pif1^4A^) impaired the function of the Pif1 NLS, as judged by the reduction of nuclear accumulation and tolerance to overexpression of these mutants. To confirm that this region contains a fully functional NLS, we fused the Pif1 775-803 region to eGFP and confirmed that it was sufficient to drive its nuclear accumulation (Fig. 4e). Consistently, this was impaired by mutation to alanines in the first basic patch. These results confirm the existence of a C-terminally located NLS in Pif1 and that the ^781^KKRK^784^ basic patch is essential for the nuclear import of Pif1, in line with a recent report^31^.

**Fig 4.**
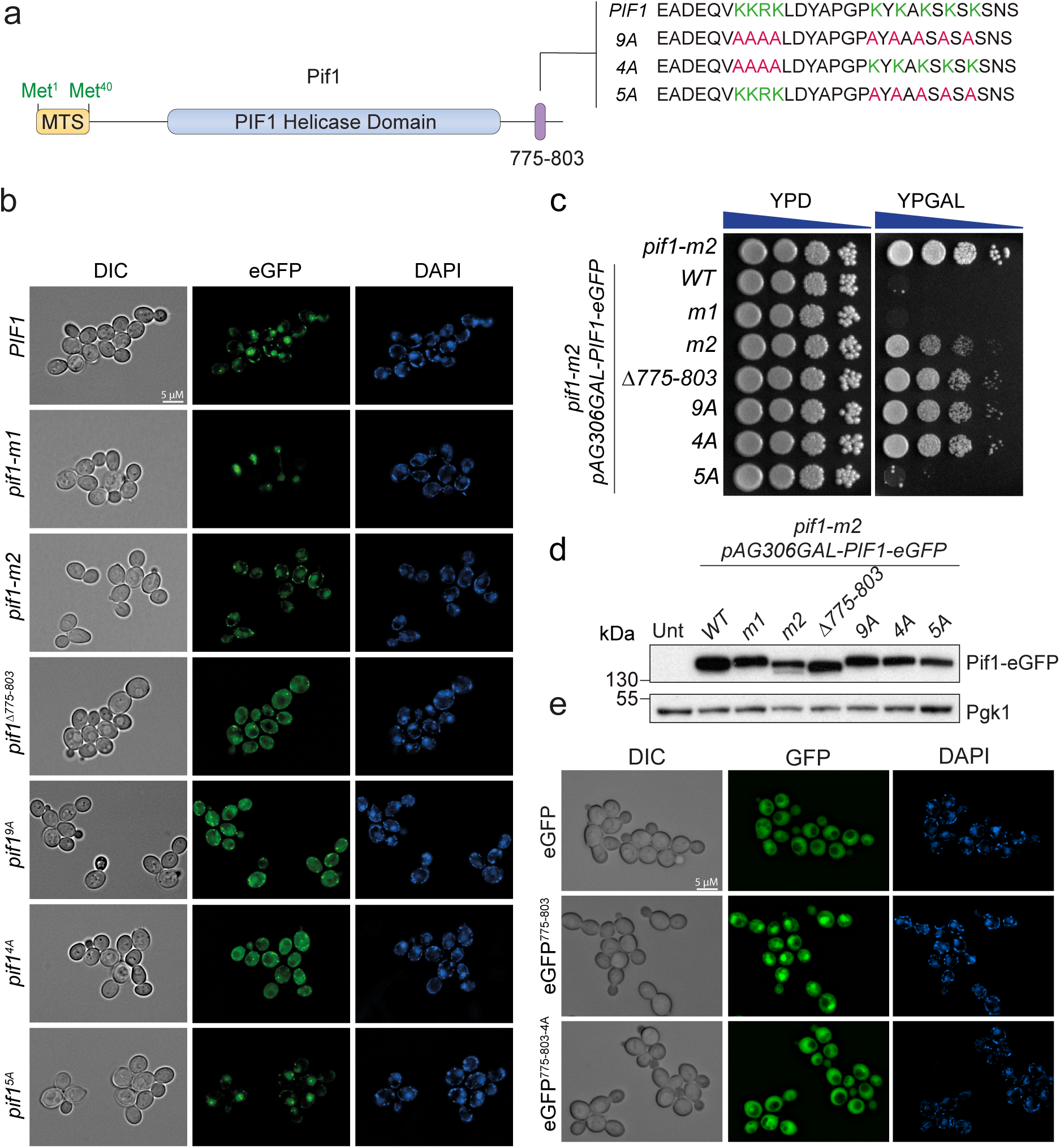
The basic patch ^781^KKRK^784^ is required for Pif1 nuclear import. (a) Schematic representation of the 775-803 region of Pif1 (purple), highlighting the amino acids substituted to alanine (green/red) in the indicated mutants. (b) Disruption of the ^781^KKRK^784^ basic patch abolishes Pif1 nuclear import. Subcellular localization of the indicated Pif1 variants in live yeast cells was assessed by fluorescence microscopy as in Figure 3a. (c) Disruption of the ^781^KKRK^784^ basic patch prevents the lethality of Pif1 overexpression. Viability of the indicated strains upon overexpression of the indicated Pif1 variants was assessed as in Figure 3c. (d) Expression levels of various Pif1-eGFP NLS mutants is comparable. Abundance of Pif1 variants in the strains from (c) was compared by western blotting using anti-GFP antibodies. Protein extracts were prepared from liquid cultures grown in YPRaff and induced for 3 h by addition of 2% galactose. Pgk1 was used as a loading control. (e) Fusion of a wild-type 775-803 Pif1 fragment to eGFP is sufficient to drive its nuclear accumulation. The subcellular localization of the indicated eGFP-fusions was assessed as in Figure 3a.

### A nuclear-null allele of Pif1 disambiguates unclear genetic interactions

Following the identification of Pif1 NLS, we decided the assess the consequences of impaired nuclear import for Pif1. For this, we employed the *pif1^4A^* allele (Fig. 4a)., which hereinafter we will rename as *pif1^nls^* (this is to avoid nomenclature overlapping with the previously described *pif1*^T763A,S765A,S766A,S769A^ phosphomutant allele^6^, also referred to as *pif1^4A^*). First, to exclude if the substitutions in Pif1^nls^ could affect its catalytical properties, we purified Pif1^nls^ as described for Pif1 (Supplementary Fig. 3c) and compared its ATPase and helicase activities to the wild-type protein (Supplementary Fig. 5a-d). In both cases, Pif1^nls^ displayed similar reaction kinetics to Pif1, indicating that the four mutations in the NLS do not interfere with its enzymatic activity.

We then tested the effect of NLS mutations on Pif1 genetic interactions by combining *pif1^nls^* with the *dia2Δ*, *dna2Δ* and *top3Δ* mutants followed by tetrad dissection analyses. Disruption of Pif1 NLS resulted in weak, but detectable, synthetic sickness with the *dia2Δ* mutants (Supplementary Fig. 5e). However, *pif1^nls^* was able to both suppress *dna2Δ* lethality (Supplementary Fig. 5f) and induce lethality in the *top3Δ* mutants (Supplementary Fig. 5g). These results indicate that disruption of the NLS in an otherwise wild-type *PIF1* allele severely compromises its nuclear functions.

Despite the NLS inactivation in *pif1^nls^*, we were still surprised that this allele did not fully recapitulate the *pif1Δ* effects on the growth of *dia2Δ* and *dna2Δ* mutants (Supplementary Fig. 5b and c). We hypothesized that even in *pif1^nls^*strains, some fraction of functional Pif1 can reach the nucleus. To minimize even further this residual nuclear Pif1 activity, we combined the mutations in both the *pif1^nls^* and *pif1-m2* alleles into a new allele which we have termed *pif1^mit^* (Fig. 5a) due to its improved separation-of-function features (Fig. 5b-d). Tetrad dissection analyses revealed that when combined with *dia2Δ* mutants, *pif1^mit^* phenocopies *pif1Δ* synthetic sickness more closely than *pif1-m2* (Fig. 5b). Likewise, the suppression of *dna2Δ* lethality by *pif1^mit^* is similar to that of *pif1Δ*, and more robust than for *pif1-m2* mutants (Fig. 5c). Moreover, *pif1^mit^* strains also display slightly increase telomeric length compared to *pif1-m2* mutants (Fig. 5d). These results indicate that *pif1^mit^* represents an improved separation-of-function allele that translates almost exclusively into mPif1.

**Fig 5.**
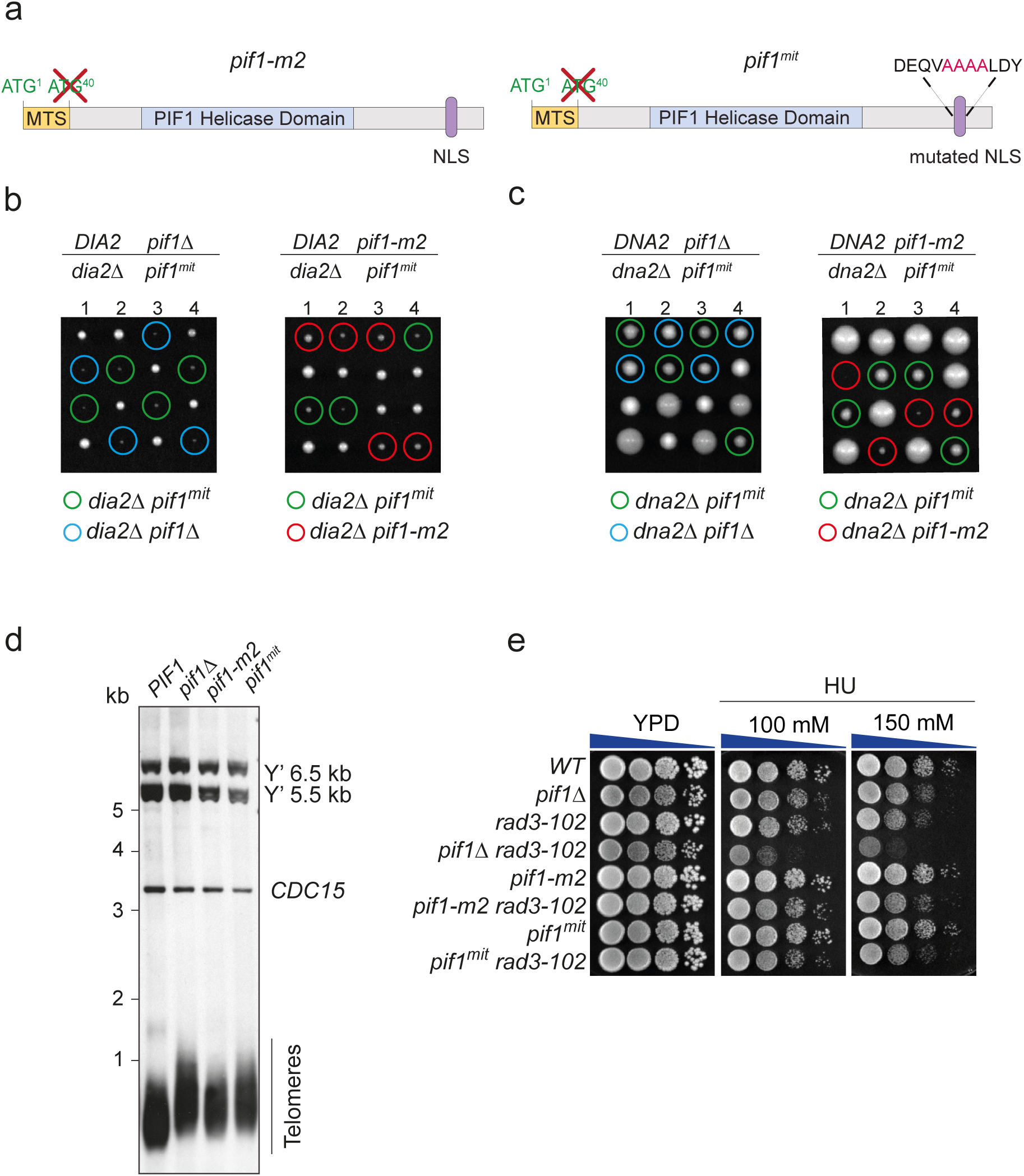
*pif1^mit^*constitutes an improved separation-of-function allele with respect to *pif1-m2*. (a) Schematic comparison of the *pif1-m2* and *pif1^mit^* alleles. (b) *pif1^mit^* recapitulates the synthetic sickness of *pif1Δ* in *dia2Δ* strains. Tetrad microdissections of the indicated diploids strains were carried out as in Figure 1a. (c) *pif1^mit^* suppresses the lethality of *dna2Δ* strains similarly to *pif1Δ*. Tetrad microdissections of the indicated diploids strains were carried out as in Figure 1c. (d) Strains expressing *pif1^mit^* display extended telomeres comparable to those in *pif1Δ* strains. Telomeric Southern blots of the indicated strains were carried out as in Figure 1g. (e) *pif1^mit^* does not increase the hypersensitivity to HU of *rad3-102* mutants. Sensitivity assays were carried out as in Figure 1b.

Interestingly, *pif1^mit^* expression in *rad3-102* cells did not increase their sensitivity to HU with respect to *PIF1* or *pif1-m2* (Fig. 5e). Since these *PIF1* alleles retain mitochondrial Pif1, this result underscores the possibility that the hypersensitivity of *pif1Δ rad3-102* mutants to HU arises from the loss of mitochondrial Pif1 activity. However, we did not detect any increased sensitivity to HU when we analysed the *pif1-m1 rad3-102* mutants (Fig. 1b), which should be devoid of mitochondrial Pif1. Hence, it cannot be ruled out that, akin to *pif1-m2*, the separation of function provided by the *pif1-m1* allele may also be incomplete.

### Residual mitochondrial Pif1 is present in *pif1-m1* mutants

In addition to the aforementioned result (Fig. 5e), during the course of our studies, we also observed an anomalous behaviour of the *pif1-m1* strains (Fig. 6a) with respect to their expected loss of mitochondrial function. While haploid *pif1Δ* strains freshly generated by tetrad dissection of *PIF1/pif1Δ* diploids had acquired an overt *petite* phenotype by the time the colony is visible, *pif1-m1* colonies arising from *PIF1/pif1-m1* diploids had not. To formally confirm this observation, we created diploid *PIF1/pif1Δ* strains with one additional ectopic copy of the *pif1-m1* and analysed their meiotic progeny by tetrad dissection. *pif1Δ* colonies displayed a clear *petite* phenotype as judged by their lack of pigmentation and inability to grow in a medium lacking a fermentable carbon source (Fig. 6b and c). Contrarily, when *pif1Δ* co-segregated with the ectopic *pif1-m1* allele, normal pigmentation and the ability to grow on YPEGly were restored (Fig. 6b and c), suggesting that this theoretically mitochondrial-null allele may also be subject to some sort of genetic bypass.

**Figure 6.**
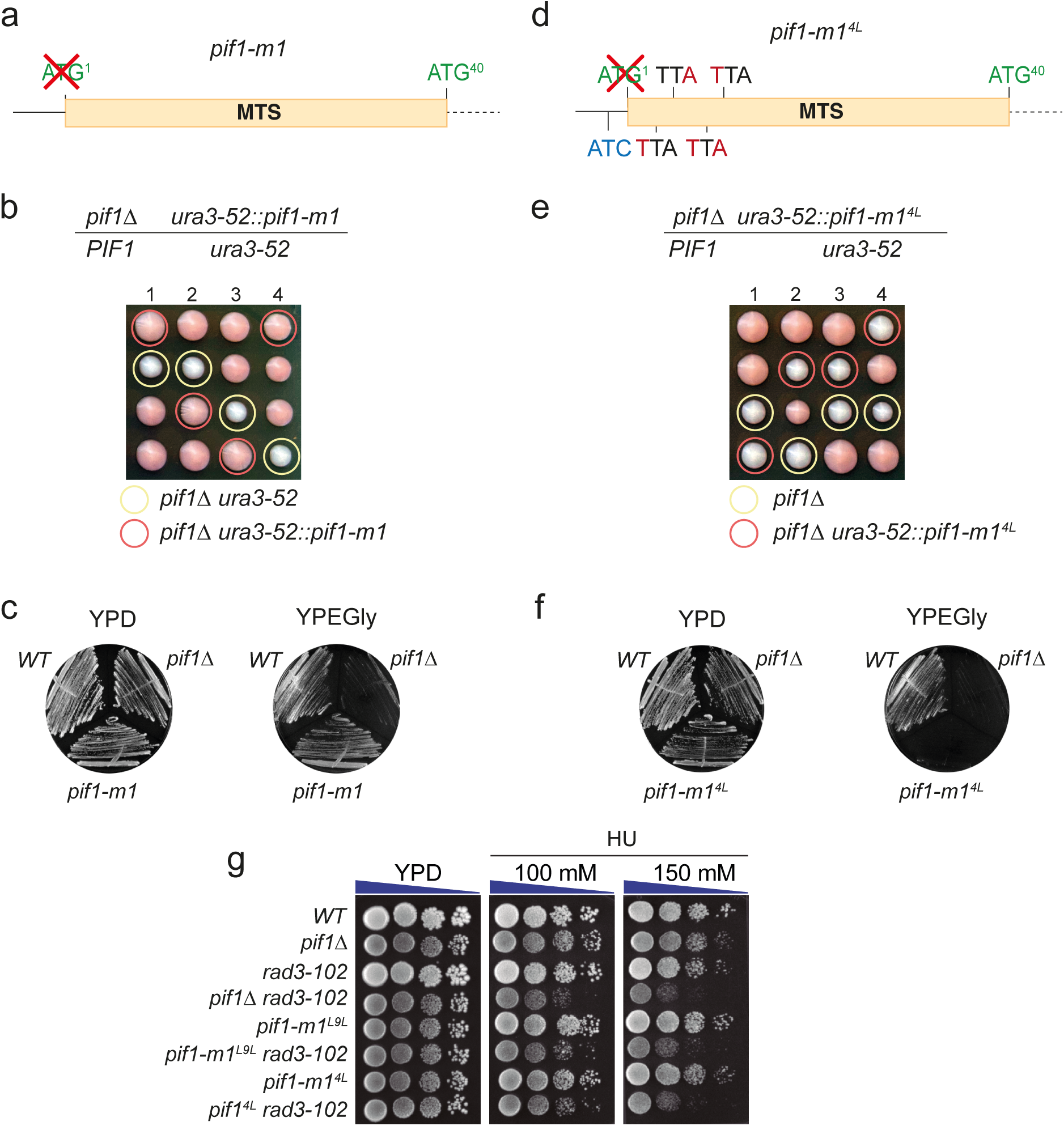
ATI from near-cognate start codons underlies incomplete separation-of-function of the *pif1-m1* allele. (a) Schematic representation of the N-terminal MTS region of *pif1-m1*. Start codons are depicted in green. (b) *pif1-m1* strains do not display a *petite* phenotype after tetrad dissection. Tetrads from a diploid strain carrying a heterozygous deletion of *PIF1* and an ectopic *pif1-m1* allele integrated at *ura3-52* were microdissected on YPD plates and incubated at 30 °C for 3 days. Adenine was omitted from YPD to allow for colour discrimination of *petite* colonies. (c) Non-fermentable carbon sources can sustain growth of *pif1-m1* strains. Freshly generated strains harbouring the indicated *PIF1* alleles were streaked on rich medium with a fermentable (YPD, 2% glucose) or non-fermentable (YPEGly, 3% ethanol, 3% glycerol) carbon source. Plates were incubated at 30 °C for 3 days and photographed. (d) Schematic representation of the N-terminal MTS region of the *pif1-m1^4L^*, depicting start codons (green), as well as the mutations in downstream near-cognate ATGs (red). (e) *pif1-m1^4L^* strains display a *petite* phenotype after tetrad dissection. As in (b), but the diploid strain harbours an ectopic copy of *pif1-m1^4L^*. (f) Non-fermentable carbon sources cannot sustain growth of *pif1-m1^4L^* strains. Strains carrying the indicated alleles were streaked as in (c). (g) *pif1-m1^4L^*, *pif1-m1^L9L^* and *pif1Δ* confer similar HU hypersensitivity to *rad3-102* mutants. Ten-fold serial dilutions of strains with the indicated genotypes were plated on YPD containing different concentrations of hydroxyurea (HU) and imaged after 2 days of incubation at 30 °C.

While this situation could seem reminiscent of the ATI by ribosomal leaky scanning observed for *pif1-m2*, there are no additional in-frame start codons in the vicinity of AUG^1^ that could support the translation initiation of a protein with a functional MTS. In this sense, it is well established that ribosomes can initiate translation at near-cognate start codons when they are embedded in a good Kozak context^32,33^. We identified five in-frame near-cognate codons in the region of AUG^1^ (one upstream: AUC^-5^ (Ile); four downstream: AUA^5^ (Ile), UUG^9^ (Leu), AUU^12^ (Ile), AUA^13^ (Ile)) that could give rise to Pif1 isoforms with functional MTSs (Suppl. Fig. 6a and b). Therefore, we introduced a series of mutations in these near-cognate codons in a *pif1-m1* construct, being as conservative as possible with respect to amino acid changes in those downstream AUG^1^ (Supplementary. Fig 6c). While mutation of the upstream near-cognate codon (*pif1-m1*^I(–5)*A*^) had a minor effect on the mitochondrial proficiency of *pif1-m1* strains (Supplementary Fig. 7a-c), substituting the four downstream codons to different leucine codons (*pif1-m1^4L^*) resulted in a complete loss of Pif1 mitochondrial function (Fig. 6d-f), with most of this effect arising from the single substitution of codon UUG^9^ (*pif1-m1^L9L^*) (Supplementary Fig. 7d-f). Importantly, the mitochondrial defects in *pif1-m1^4L^*are not due to reduced protein levels (Supplementary. Fig 6d) or disruption of Pif1 MTS, as the same substitutions in a wild-type allele (*pif1^4L^*) do not result in a *petite* phenotype (Supplementary Fig. 7g-i). Collectively, these results indicate that translation initiation from near-cognate start codons in the vicinity of AUG^1^ can give rise to biologically relevant amounts of mitochondrial Pif1, which, in our hands, can suppress the mitochondrial defects predicted for *pif1-m1*.

With the *pif1-m1^4L^* allele in hand, we decided to test whether the hypersensitivity of *pif1Δ rad3-102* mutants to HU derives from the loss of mitochondrial Pif1. In agreement with this hypothesis, the presence of the *pif1-m1^4L^ or pif1-m1^L9L^* in *rad3-102* mutants resulted in similar HU hypersensitivity to the *pif1Δ rad3-102* strain (Fig. 6h), revealing a counterintuitive mitochondrial connection between Pif1 and Rad3 in their response to replication stress. These results underscore the importance of the development of improved separation-of-function alleles in order to disambiguate the biological roles of this helicase in the nucleus and mitochondria.

## Discussion

In this work, we have determined that ribosomal leaky scanning is the specific ATI mechanism responsible for the generation of mPif1 and nPif1. Under this model, the stringency or weakness of the Kozak context influences the probability for the scanning 40S subunits to engage or skip the first AUG, which could enable translation initiation from downstream AUG codons. The sub-optimal Kozak context surrounding AUG^1^ in *PIF1* mRNA (Supplementary Fig. 2f) allows a significant fraction of the scanning ribosomes to bypass AUG^1^, responsible for the translation of mPif1, and start from AUG^40^, resulting in the translation of nPif1. However, our results demonstrate that the weak Kozak motif around AUG^40^ (Supplementary Fig. 2f) is permissive enough to let a minor fraction of scanning ribosomes to traverse until the next in-frame start codon, AUG^107^, and produce detectable amounts of the shorter Pif1^107–859^ isoform even in wild-type cells (Fig. 2b). Consistently with this leaky scanning model, mutations in the preceding start codon (AUG^40^), as in *pif1-m2* mutants, enable a higher proportion of ribosomes to reach AUG^107^ and increase the abundance of this isoform (Fig. 2b and c), while an enhanced engagement of ribosomes by AUG^40^ renders this isoform undetectable (Fig. 2f). Moreover, the sole mutation of AUG^107^ is not sufficient to abrogate the generation of a fast-migrating isoform. Instead, the complete disappearance of this band requires the concomitant mutation of the AUG^107^/AUG^113^/AUG^127^ cluster, further substantiating the intrinsic tendency of *PIF1* mRNA to undergo ATI by leaky scanning (Fig. 2e).

Under this perspective, we argue that the incomplete separation of function observed for the *pif1-m1* and *pif1-m2* alleles is a consequence, at least partially, of this tendency of *PIF1* mRNA to undergo ribosomal leaky scanning (Fig. 7). In the case of *pif1-m1* mutants, the elimination of AUG^1^ is not sufficient to produce an overt mitochondrial phenotype, suggesting the presence of relevant amounts of mPif1. In agreement with the absence of any in-frame AUG codons that would enable the translation of MTS-containing isoforms, we demonstrated that near-cognate start codons in the vicinity of AUG^1^ may serve as alternative translation initiation points (Fig. 6 and 7). Usage of these non-AUG codons would result in isoforms that still contain a functional MTS, rendering mature mPif1 (Supplementary Fig. S6b). Despite being undetectable by western blot, such mPif1 supports mitochondrial function in *pif1-m1* mutants, allowing their growth on non-fermentable carbon sources (Fig. 6c). In *pif1-m2* mutants, the increased amounts of the biochemically proficient and nuclear Pif1^107–859^ isoform provides a plausible explanation for the oft-invoked residual nuclear function of these mutants ^11–17^, as suggested by the similar restoration of cell growth produced by the expression of *pif1*^107–859^ or *PIF1* in *dia2Δ* mutants (Fig. 3d). However, our results are more compatible with a scenario where the residual nuclear activity in *pif1-m2* mutants derives from a combined effect between the increased levels of Pif1^107–859^ and, akin to *pif1-m1* mutants, the existence of other undetectable isoforms produced by translation initiation from non-AUG codons (Fig. 7). This is supported by the existence of several near-cognate codons with a favourable context near AUG^40^ that display an enrichment in scanning small ribosomal subunits by TCP-seq (Supplementary Fig. 8) and by the reduced functionality of *pif1*^107–859^ compared to *PIF1* in two different scenarios: i) *pif1*^107–859^ overexpression does not impair cell growth (Fig. 3a) and ii) *dna2Δ* cells maintain viability in the presence of *pif1*^107–859^ (Fig. 3d). Noteworthy, the Pif1 N-terminal domain (40-237) is responsible for the toxicity associated to *PIF1* overexpression ^34^ and it is the target of several PTMs, including checkpoint- and cell-cycle dependent phosphorylation^7,35^, as well as acetylation^36^. Then, it is arguable that Pif1^107–859^ might lack regulatory domains located between amino acids 40-107 that could be included in isoforms arising from non-AUG codons within that region, thus allowing them to fulfil those functions that Pif1^107–859^ cannot satisfy.

**Figure 7.**
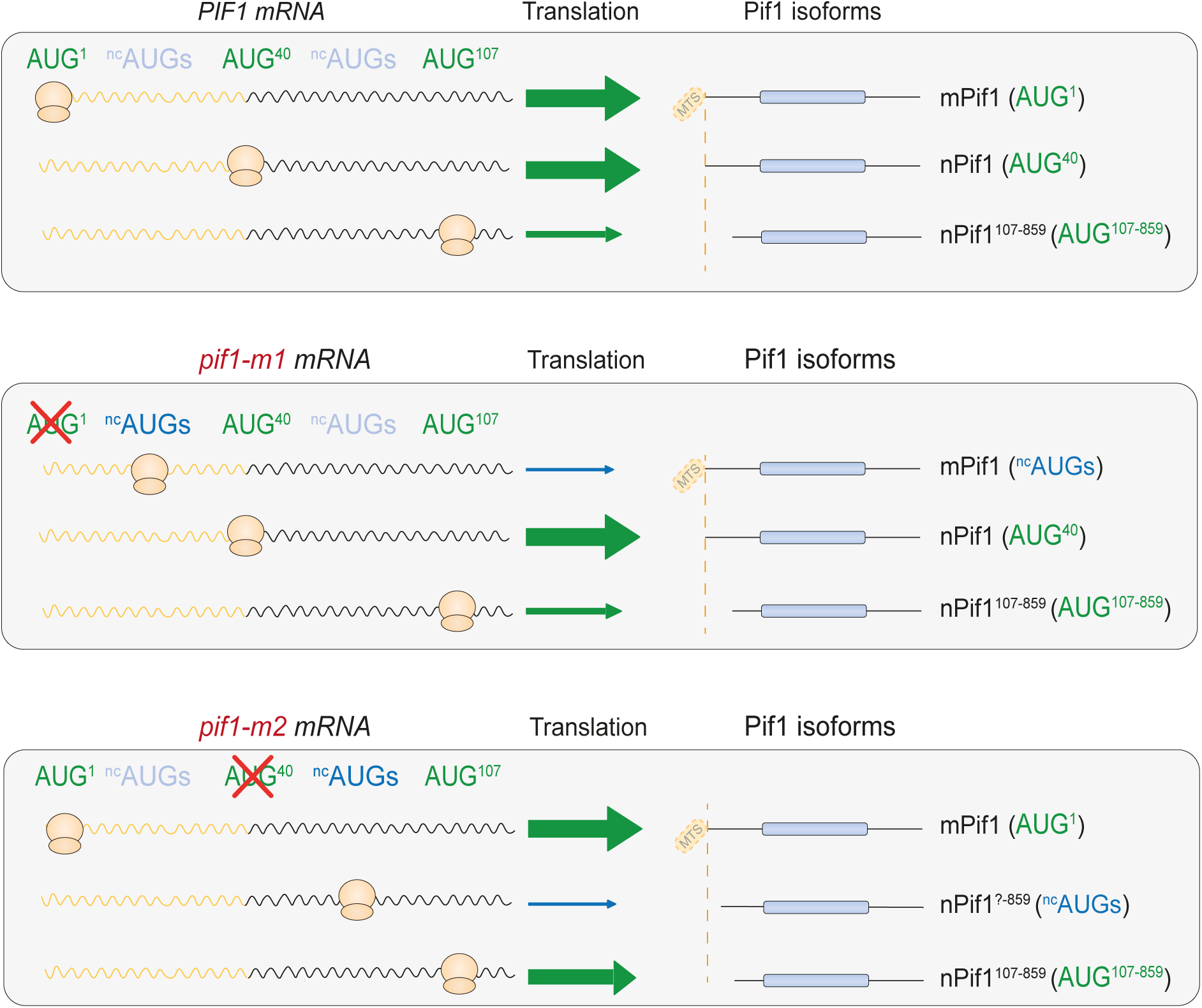
A refined model for the translation of *PIF1* mRNA. *PIF1* mRNA (upper panel) undergoes ATI from at least three AUG codons (green). AUG^1^ and AUG^40^ are responsible for the production of the main isoforms of mPif1 and nPif1, with a small fraction of ribosomes reaching AUG^107^ to generate the nuclear Pif1^107-859^. Minor contributions from near-cognate start codons (^nc^AUGs, blue) are not depicted here. In *pif1-m1* mutants (middle panel), mutation of AUG^1^ increases translation from proximal, downstream ^nc^AUGs. Despite their small N-terminal truncations, these mPif1 precursors retain an operative MTS and therefore may sustain mitochondrial function. In *pif1-m2* mutants (bottom panel), the absence of AUG^40^ allows a higher proportion of scanning ribosomes to reach AUG^107^ and increase the production of nPif1^107-859^, which may support some of nPif1. The potential contribution of ^nc^AUGs between AUG^40^ and AUG^107^ to produce additional isoforms of nPif1 with intermediate truncations is also indicated.

Considering all this plasticity for translation initiation of *PIF1* mRNA, we have built on the *pif1-m1* and *pif1-m2* alleles to minimize their residual mitochondrial and nuclear functions, respectively. In this sense, a recent report has employed a similar rationale to us in order to minimize the nuclear levels of Pif1 by independently identifying and disrupting the so-far uncharacterized Pif1 NLS^31^. Both works identify the ^781^KKRK^784^ basic patch as required for Pif1 nuclear functions, while dispensable for its mitochondrial roles, by either deleting these residues^31^ or mutating them to alanines (Fig. 5 and Supplementary Fig. 5). The involvement of this sequence in nuclear import was also independently confirmed either by suppression of the *pif1-NLSΔ* mutant phenotypes by fusing it to an SV40 NLS^31^ or directly by fluorescence microscopy, showing: i) loss of Pif1-eGFP nuclear accumulation when this patch is mutated (Fig. 4b) and ii) nuclear accumulation of eGFP when fused to the Pif1 fragment containing this basic patch (Fig. 4e). Moreover, our results also confirm that nuclear phenotypes in an NLS-deficient *PIF1* mutant do not arise reduced catalytic activity of the helicase (Supplementary Fig. 5). The impairment of Pif1 NLS does not suffice to completely abolish Pif1 nuclear functions (Supplementary Fig. 5). Potential reasons behind this observation, including diffusion through the nuclear pore, a secondary NLS in Pif1 or association to a carrier protein, have been elegantly laid out by Lee et al.^31^ and will not be further discussed here. Given these results, both groups have resorted to the impairment of the NLS in *pif1-m2* to further diminish nuclear Pif1 levels. Therefore, both *pif1-m2-NLSΔ*^31^ and *pif1^mit^*(Fig. 5) represent the best alleles to date to abrogate nuclear Pif1 functions without affecting its mitochondrial activities. Additionally, we have combined mutations in near-cognate start codons with *pif1-m1* to produce the improved mitochondrial-deficient allele *pif1-m1^4L^* (Fig. 6). We envisage that the combination of *pif1^mit^* and *pif1-m1^4L^* (= *pif1^nuc^*) will enable the discovery of subtle genetic interactions that could have been previously missed due to the residual activities of *pif1-m1* and *pif1-m2* or the confounding effect of *pif1Δ* pleiotropy. As a proof of principle, we have disambiguated the origin of the hypersensitivity of *pif1Δ rad3-102* mutants to HU^37^, which, unexpectedly, arises from the loss of Pif1 mitochondrial activity, rather than nuclear (Fig. 6g). Therefore, these alleles may contribute to establish more controlled genetic set-ups for the study of Pif1 in the replication and repair of the mitochondrial and nuclear genomes.

This plasticity in the translation initiation step has important implications for the design of separation-of-function alleles that are likely applicable to other dual-targeted DNA repair factors or, more generally, to any other genes where ATI is responsible for the generation of functionally different isoforms. Akin to the problematic described for *S. cerevisiae,* in *S. pombe*, the *PIF1* ortholog *pfh1* also generates essential nuclear and mitochondrial isoforms through ATI, but nuclear-deficient alleles of *pfh1* (mutated for ATG^21^, the start codon for nuclear Pfh1) can sustain cell division indefinitely (Pinter et al., 2008), suggesting an additional source of nuclear Phf1. In fact, the combined mutation of ATG^21^ and the next three in-frame, downstream ATGs, does not suffice to recapitulate the *pfh1^-^* phenotype^38^. This strongly suggests that near-cognate start codons, like in *PIF1*, could also enable ATI of nuclear Pfh1 isoforms that become phenotypically relevant in the absence of the main start codons^39^

It is intuitive to expand the potential impact of ATI for the study of other dual-targeted genes in budding and fission yeasts. For instance, the conserved ligase I (*CDC9* in *S. cerevisiae* / *cdc17* in *S. pombe*) also presents nuclear and mitochondrial isoforms derived from ATI at two in-frame start codons flanking its MTS^40^. The phenotypes exhibited by their separation-of-function alleles again suggest that individual mutations of the first and second start codons do not completely abolish the production of mitochondrial or nuclear isoforms. In budding yeast, mutation of the *CDC9* second ATG, which should abrogate the expression of the nuclear isoform, still yields viable cells and, like in *pif1-m2*, correlates with the appearance of a fast-migrating isoform^40^. In *S. pombe*, the equivalent mutation of the second ATG (ATG^20^) in *cdc17* failed to completely abolish nuclear function unless the NLS was also mutated^41^. Given that no in-frame start codons exist before this NLS, the most likely explanation for the residual nuclear protein upon mutation of ATG^20^ would be ATI events from two near-cognates at positions 27 and 32^41^.

In human cells, ATI by ribosomal leaky scanning and near-cognate start codon usage also contribute to the generation of protein variants, frequently in combination with alternative splicing, thus leading to similar scenarios to that of Pif1 in budding yeast. For instance, the mRNAs of disease-linked DNA repair factors TOP3α^42^ and RNAseH1^43^ undergo ribosomal leaky scanning for their dual targeting to nucleus and mitochondria. Both genes present downstream ATGs and near-cognates with strong Kozak contexts that could sustain a relevant amount of translation initiation in mutants of the primary start codons. Consistently, in order to achieve the exclusive expression of the mitochondrial isoform of human TOP3α, both the mutation of both its second ATG (which drives the translation of the nuclear isoform) and its NLS is required^44^. Incomplete separation-of-function mutants for dual-targeted DNA repair factors have been also described for human PIF1^45^ and ADAR1^46,47^. Recently, the striking difference between the complete inactivation of the APC/C complex coactivator CDC20 and its first ATG codon has also been highlighted. Unlike the full-length protein, the N-terminally truncated isoforms produced by ribosomal leaky scanning that specifically increase after mutation of the first *CDC20* ATG codon cannot be inhibited by the spindle-assembly checkpoint and therefore allow progression into mitosis despite SAC activation^48^.

Cumulative evidence from genome-wide analyses and bioinformatic predictions indicate that thousands of genes may undergo ATI events from both in-frame AUG and near-cognate start codons. This enriches the diversity of the cellular proteome by producing extended or truncated protein isoforms, with the potential gain or loss of functional domains in physiological and pathological situations^49–57^. Given this complexity, genetic approaches mutating the main start codons to understand the function of specific protein isoforms must consider the potential contribution of ATI events to the protein pool.

## Acknowledgments

The Blanco lab was supported from MCIN / AEI / 10.13039/501100011033 [PID2020-115472GB-I00]; Xunta de Galicia (XdG) / FEDER ‘Una manera de hacer Europa’ [ED431C 2019/013]; CIMUS receives financial support from the XdG / FEDER [ED431G 2019/02, Centro Singular de Investigación de Galicia, accreditation 2019–2022]; T.L.-D. was a recipient of a pre-doctoral fellowships from XdG [ED481A-2018/042]. The authors would like to thank Virginia Zakian and Rodrigo Bermejo for various yeast strains, Joao Matos and David Lydall for plasmids, and Maria Crugeiras for the critical reading of the manuscript.

## Author contributions

T.L.-D. carried out all the experimental work. T.L.-D. and M.G.B. conceived the project, designed experiments, analyzed data and wrote the manuscript.

## Methods

### Yeast manipulation and growing

All *S. cerevisiae* strains are derivatives of W303 background, as detailed in Supplementary Table 1. Yeast manipulations were performed according to standard procedures^58^. Cells were typically grown in YP (1% yeast extract, 2% peptone) containing 2% glucose (YPD) or 2% raffinose (YPRaff). For induction of genes under the control of the *GAL1* promoter, 2% galactose was added to YPRaff (YPGal). YP containing 3% ethanol and 3% glycerol (YPEGly) was employed to test *petite* phenotypes. SC synthetic media were employed for selection of auxotrophic mutants and for live cell microscopy. In some cases, adenine (40 µg/ml) was omitted from media to allow development of the red pigment in W303 strains. Mutant strains were generated by PCR-based editing^59,60^, followed by diploid and tetrad dissections when required. Point mutations at endogenous loci were introduced with *delitto perfetto*^61^.

### Tetrad dissection analysis

Strains of opposite mating types were mixed in patches and incubated on YPD plates for 4 h at 30°C. Diploids were selected by consecutive streaks in selective media, or by micromanipulation under a dissection microscope. Once selected, diploids were incubated on SPO plates (CH_3_COOK 3 g/l, 10 µg/ml Ade, 10 µg/ml His, 10 µg/ml Leu, 10 µg/ml Trp, 10 µg/ml Ura) for 3 days at 30 °C before microdissection. After dissections, plates were incubated at 30 °C and imaged after 48, 72 and 96 h in a Gel Doc XR+ (Bio-Rad).

### Cloning and mutagenesis

All plasmids in this study are listed in Supplementary Table 2. The coding sequences of C-terminal 6xHis-6xFLAG-tagged *PIF1*, *pif1-m1* and *pif1-m2* were cloned in pRSII406 vectors^62^. Alternatively, untagged versions of the same alleles were cloned in pENTR221 and shuttled to pAG306GAL-ccdB-eGFP vectors^63^. From these vectors, derivative plasmids carrying different mutations or truncations were created by standard subcloning, inverse PCR and/or Gateway recombination. Details of each cloning can be provided upon request.

### Protein analysis

Yeast cultures in mid log-phase were harvested and cell pellets disrupted using glass beads in 10% TCA. Precipitates were collected by centrifugation, resuspended in 2x NuPAGE sample buffer (Invitrogen, #NP0008) supplemented with 200 mM DTT, and neutralized with 1 M Tris-base. Samples were boiled at 95°C for 5 min, cleared by centrifugation, and separated in either NuPAGE 3-8% (Invitrogen, #EA0375BOX) or 7% Tris-Acetate polyacrylamide gradient gels (Invitrogen, #EA0355BOX).

For Western blotting, proteins were transferred onto Amersham™ Hybond® P PVDF membranes (GE Healthcare, #10600023) and detected with the following antibodies: FLAG-HRP (mouse, 1:10000, A8592-1MG, Sigma), eGFP (mouse 1:2000, 11814460001, Roche) and Pgk1-HRP (mouse, 1:5000, ab197960, Abcam). Alternatively, for fluorescence detection and quantitative western blotting, total proteins were detected with No-Stain Reagent (Invitrogen, #A44449) after transfer onto Immobilon - FL PVDF membranes (Millipore, #IPFL00010). FLAG-tagged proteins were detected using M2 anti-FLAG antibody (mouse, F1084, Merck) followed and fluorescent Anti-Mouse Alexa Fluor™ Plus 800, (Goat, 1:10000, A32730, Invitrogen) and imaged in a ChemiDoc™ MP (Bio-Rad).

### Protein purification

A pET28b plasmid encoding C-terminal 6xHis-tagged nuclear Pif1 (Addgene plasmid #65047)^64^ was employed to express and purify Pif1 from *E. coli* BL21-CodonPlus (DE3)-RIL strain (Agilent) as described ^65^. The same plasmid was mutagenized to express and purify Pif1^nls^, Pif1^107–859^ and ATPase-dead Pif1^KA^ (K264A), employing the same protocol.

### Splayed arm substrate preparation

Splayed arm substrates with 3’-6FAM labelling were prepared essentially as previously described^66^, employing the following PAGE-purified ssDNA oligonucleotides (Sigma-Aldrich):

A6-3’FAM (5’-ATTGGTTATTTACCGAGCTCGAATTCACTGG-3’-6FAM) and A9 (5’- CCAGTGAATTCGAGCTCGGTACCCGCTAGCGGGGATCCTCTA-3’)^67^.

Briefly, labelled and unlabelled oligonucleotides were mixed in a 1:3 ratio, boiled in a water bath and cooled down to room temperature overnight. Substrates were subsequently purified from 10% polyacrylamide gels and eluted in TMgN buffer (10 mM Tris-HCl pH 8.0, 1 mM MgCl_2_, 50 mM NaCl).

### ATP hydrolysis assays

Pif1 (10 nM) was incubated with 0.165 µM [^32^P-γ]-ATP at 30 °C in 15 µl of ATPase buffer (35 mM Tris-HCl, pH 7.5, 1mM DTT, 5 mM MgCl_2_, and 100 µg/ml BSA, 42 mM KCl) containing 1 µM poli-T ssDNA. Reactions were terminated at the indicated times transferring 2 µl to pre-aliquoted tubes with 2X Stop solution (2% SDS, 40 mM EDTA). Thin-layer chromatography (Z122882-25EA, Sigma-Aldrich® TLC plates) and phosphorimaging analysis to quantify ATP hydrolysis were conducted as described elsewhere^68^, using a Typhoon FLA 9500 scanner and the ImageQuant software (GE Healthcare).

### DNA unwinding assays

Pif1 (4 nM) was incubated with a 10-molar excess of splayed arm substrate at 30 °C for 3 min in 10 µl of buffer H (35 mM Tris-HCl, pH 7.5, 1 mM DTT, 5 mM MgCl_2_, 100 ng/µl BSA, 5 mM ATP, 60 mM KCl). A 100-molar excess of ssDNA trap (unlabelled A6 oligo, see Splayed Arm substrate preparation) was added to avoid reannealing of the fluorescent-labelled oligo. The reactions were terminated at the indicated times by deproteinization with 0.5% SDS and proteinase K (2 mg/ml) at 37 °C for 30 min. DNA species were resolved through native 12% polyacrylamide gels in TBE buffer. Gels were imaged on a Typhoon FLA 9500 and quantified with ImageQuant software (GE Healthcare)

### DNA damage sensitivity assays

Cells grown to mid-log phase were normalized to OD_600_=0.5 and 10-fold serial dilutions were spotted onto YPGal or YPDA with different concentrations of hydroxyurea (H9120, US Biological). Plates were incubated for 3 days at 30 °C and imaged in a Gel Doc XR+ (Bio-Rad).

### Telomeric Southern Blot

*PIF1/pif1Δ* diploid strains carrying an additional copy of different *PIF1* alleles, ectopically integrated in *ura3-52*, were sporulated and dissected at the same time. After replica-plating (1^st^ passage), strains with genotypes of interest were selected (2^nd^ passage) and re-streaked again (3^rd^ passage) to allow telomere elongation in Pif1-deficient strains. Starter cultures were prepared by pooling three biological replicates for each genotype. Telomere length was assayed by Southern hybridization as described ^13^. Genomic DNA was first purified by phenol-chloroform and ethanol precipitation. Subsequently, DNA was further purified by silica-columns. 3 µg of genomic DNA was digested with *XhoI* overnight. 1 µg of each sample was loaded on 1% (w/v) agarose gels (20 cm length) and separated for 18 h at 1.5 V/cm in TBE 0.5X. After electrophoresis, DNA fragments were blotted onto Zeta-Probe nylon membranes (Bio-Rad, #1620159) by the alkaline transfer method, according to manufacturer’s instructions. Synthesis of the labelled probes, membrane hybridization and detection were performed using DIG-High Prime DNA Labelling and Detection Starter Kit II (Roche, #11585614910) as per the manufacturer’s instructions. Membranes were imaged on a Typhoon FLA 9500. The subtelomeric Y’ probe was amplified from pDL987 (kind gift from David Lydall) using oligos OMB363 and OMB364, respectively. The *CDC15* probe was synthesized using oligos OMB349 and OMB350, as previously described^69^. The GeneRuler 1 kb Plus DNA Ladder (ThermoFisher, #SM1331) was employed as a molecular-weight marker.

### Fluorescence microscopy

Pif1 subcellular localization was addressed by fluorescence microscopy in live yeast cells as described ^70^. Briefly, mid-log phase cultures in SC-Raffinose + 100 µg/ml of adenine were induced for 3 h with 2% galactose. 30 min prior to the end of induction, DAPI was added to a final concentration of 10 µg/ml. After induction, 1 ml of cells were harvested (1500 xg) and resuspended in 100 µl of SC-Raff-Gal + 100 µg/ml adenine. 3.5 µl of the cell suspension were transferred to microscope slides with a solidified patch of 1.2% agarose. The edges of the cover slip were sealed with melted VALAP before imaging. Cells were visualized using a THUNDER Imager Tissue (Leica Microsystems) with the following parameters: refractive index 1.34 and Thunder algorithm enabled for background subtraction.

### *In silico* sequence analyses

Potential regions to encode for NLSs in Pif1 sequence were identified by employing the cNLS mapper software^71^ with the discovery threshold set to 4.0, together with NucPred^72^ and NLStradamus^73^ with their default parameters. Potential mitochondrial pre-sequences were analysed with Mitofates^74^, using fungal or metazoan parameters where applicable. For the prediction of secondary structures and disordered regions in Pif1, the PSIPRED package^75^ - including PSIPRED 4.0 and DISOPRED3- was employed with the default parameters.

## References

1. Bochman, M. L., Judge, C. P. & Zakian, V. A. The Pif1 family in prokaryotes: what are our helicases doing in your bacteria? Mol. Biol. Cell 22, 1955–1959 (2011).

2. Bochman, M. L., Sabouri, N. & Zakian, V. A. Unwinding the functions of the Pif1 family helicases. DNA Repair 9, 237–249 (2010).

3. Malone, E. G., Thompson, M. D. & Byrd, A. K. Role and Regulation of Pif1 Family Helicases at the Replication Fork. Int. J. Mol. Sci. 23, 3736 (2022).

4. Muellner, J. & Schmidt, K. H. Yeast Genome Maintenance by the Multifunctional PIF1 DNA Helicase Family. Genes 11, (2020).

5. Ononye, O. E., Sausen, C. W., Balakrishnan, L. & Bochman, M. L. Lysine acetylation regulates the activity of nuclear Pif1. J. Biol. Chem. 295, 15482–15497 (2020).

6. Makovets, S. & Blackburn, E. H. DNA damage signalling prevents deleterious telomere addition at DNA breaks. Nat. Cell Biol. 11, 1383–1386 (2009).

7. Rossi, S. E., Ajazi, A., Carotenuto, W., Foiani, M. & Giannattasio, M. Rad53-Mediated Regulation of Rrm3 and Pif1 DNA Helicases Contributes to Prevention of Aberrant Fork Transitions under Replication Stress. Cell Rep. 13, 80–92 (2015).

8. O’Rourke, T. W., Doudican, N. A., Mackereth, M. D., Doetsch, P. W. & Shadel, G. S. Mitochondrial dysfunction due to oxidative mitochondrial DNA damage is reduced through cooperative actions of diverse proteins. Mol Cell Biol 22, 4086–93 (2002).

9. Guirola, M. et al. Lack of DNA helicase Pif1 disrupts zinc and iron homoeostasis in yeast. Biochem. J. 432, 595–605 (2010).

10. Schulz, V. P. & Zakian, V. A. The saccharomyces PIF1 DNA helicase inhibits telomere elongation and de novo telomere formation. Cell 76, 145–155 (1994).

11. Buzovetsky, O. et al. Role of the Pif1-PCNA Complex in Pol delta-Dependent Strand Displacement DNA Synthesis and Break-Induced Replication. Cell Rep. 21, 1707–1714 (2017).

12. Dahan, D. et al. Pif1 is essential for efficient replisome progression through lagging strand G-quadruplex DNA secondary structures. Nucleic Acids Res. 46, 11847–11857 (2018).

13. Dewar, J. M. & Lydall, D. Pif1- and Exo1-dependent nucleases coordinate checkpoint activation following telomere uncapping. EMBO J. 29, 4020–4034 (2010).

14. Paeschke, K. et al. Pif1 family helicases suppress genome instability at G-quadruplex motifs. Nature 497, 458–462 (2013).

15. Ribeyre, C. et al. The yeast Pif1 helicase prevents genomic instability caused by. PLoS Genet. 5, e1000475 (2009).

16. Sakofsky, C. J. et al. Translesion Polymerases Drive Microhomology-Mediated Break-Induced Replication Leading to Complex Chromosomal Rearrangements. Mol. Cell 60, 860–872 (2015).

17. Stundon, J. L. & Zakian, V. A. Identification of Saccharomyces cerevisiae Genes Whose Deletion Causes Synthetic Effects in Cells with Reduced Levels of the Nuclear Pif1 DNA Helicase. G3 Bethesda Md 5, 2913–2918 (2015).

18. Pan, X. et al. A DNA Integrity Network in the Yeast Saccharomyces cerevisiae. Cell 124, 1069–1081 (2006).

19. Stundon, J. L. & Zakian, V. A. Identification of Saccharomyces cerevisiae Genes Whose Deletion Causes Synthetic Effects in Cells with Reduced Levels of the Nuclear Pif1 DNA Helicase. G3 Bethesda Md 5, 2913–2918 (2015).

20. Moriel-Carretero, M. & Aguilera, A. A postincision-deficient TFIIH causes replication fork breakage and uncovers alternative Rad51- or Pol32-mediated restart mechanisms. Mol. Cell 37, 690–701 (2010).

21. Budd, M. E., Reis, C. C., Smith, S., Myung, K. & Campbell, J. L. Evidence suggesting that Pif1 helicase functions in DNA replication with the DNA2 helicase/nuclease and DNA polymerase delta. Mol Cell Biol 26, 2490–2500 (2006).

22. Wagner, M., Price, G. & Rothstein, R. The absence of Top3 reveals an interaction between the Sgs1 and Pif1 DNA Helicases in Saccharomyces cerevisiae. Genetics 174, 555–573 (2006).

23. Zhou, J.-Q. Pif1p Helicase, a Catalytic Inhibitor of Telomerase in Yeast. Science 289, 771–774 (2000).

24. Lahaye, A., Leterme, S. & Foury, F. PIF1 DNA helicase from Saccharomyces cerevisiae. Biochemical characterization of the enzyme. J. Biol. Chem. 268, 26155–26161 (1993).

25. Sriram, A., Bohlen, J. & Teleman, A. A. Translation acrobatics: how cancer cells exploit alternate modes of translational initiation. EMBO Rep. 19, e45947 (2018).

26. Hinnebusch, A. G. Molecular mechanism of scanning and start codon selection in eukaryotes. Microbiol. Mol. Biol. Rev. MMBR 75, 434–467, first page of table of contents (2011).

27. Kozak, M. Pushing the limits of the scanning mechanism for initiation of translation. Gene 299, 1–34 (2002).

28. Lahaye, A., Stahl, H., Thines-Sempoux, D. & Foury, F. PIF1: a DNA helicase in yeast mitochondria. EMBO J. 10, 997–1007 (1991).

29. Chang, M. et al. Telomerase Is Essential to Alleviate Pif1-Induced Replication Stress at Telomeres. Genetics 183, 779–791 (2009).

30. Lahaye, A., Leterme, S. & Foury, F. PIF1 DNA helicase from Saccharomyces cerevisiae. Biochemical characterization of the enzyme. J. Biol. Chem. 268, 26155–26161 (1993).

31. Lee, R. S. et al. Identification of the nuclear localization signal in the Saccharomyces cerevisiae Pif1 DNA helicase. PLOS Genet. 19, e1010853 (2023).

32. Chen, S.-J., Lin, G., Chang, K.-J., Yeh, L.-S. & Wang, C.-C. Translational Efficiency of a Non-AUG Initiation Codon Is Significantly Affected by Its Sequence Context in Yeast *. J. Biol. Chem. 283, 3173–3180 (2008).

33. Kearse, M. G. & Wilusz, J. E. Non-AUG translation: a new start for protein synthesis in eukaryotes. Genes Dev. 31, 1717–1731 (2017).

34. Nickens, D. G., Sausen, C. W. & Bochman, M. L. The Biochemical Activities of the Saccharomyces cerevisiae Pif1 Helicase Are Regulated by Its N-Terminal Domain. Genes 10, (2019).

35. Holt, L. J. et al. Global Analysis of Cdk1 Substrate Phosphorylation Sites Provides Insights into Evolution. Science 325, 1682–1686 (2009).

36. Ononye, O. E., Sausen, C. W., Balakrishnan, L. & Bochman, M. L. Lysine acetylation regulates the activity of nuclear Pif1. J Biol Chem 295, 15482–15497 (2020).

37. Moriel-Carretero, M. & Aguilera, A. A Postincision-Deficient TFIIH Causes Replication Fork Breakage and Uncovers Alternative Rad51- or Pol32-Mediated Restart Mechanisms. Mol. Cell 37, 690–701 (2010).

38. Pinter, S. F., Aubert, S. D. & Zakian, V. A. The Schizosaccharomyces pombe Pfh1p DNA Helicase Is Essential for the Maintenance of Nuclear and Mitochondrial DNA. Mol. Cell. Biol. 28, 6594–6608 (2008).

39. Shimada, K. & Gasser, S. M. DNA replication: Pif1 pulls the plug on stalled replication forks. Curr. Biol. CB 22, R404–405 (2012).

40. Willer, M., Rainey, M., Pullen, T. & Stirling, C. J. The yeast CDC9 gene encodes both a nuclear and a mitochondrial form of DNA ligase I. Curr. Biol. 9, 1085–S1 (1999).

41. Martin, I. V. & MacNeill, S. A. Functional analysis of subcellular localization and protein– protein interaction sequences in the essential DNA ligase I protein of fission yeast. Nucleic Acids Res. 32, 632–642 (2004).

42. Wang, Y., Lyu, Y. L. & Wang, J. C. Dual localization of human DNA topoisomerase IIIα to mitochondria and nucleus. Proc. Natl. Acad. Sci. 99, 12114–12119 (2002).

43. Suzuki, Y. et al. An Upstream Open Reading Frame and the Context of the Two AUG Codons Affect the Abundance of Mitochondrial and Nuclear RNase H1. Mol. Cell. Biol. 30, 5123–5134 (2010).

44. Hangas, A. et al. Top3α is the replicative topoisomerase in mitochondrial DNA replication. Nucleic Acids Res. 50, 8733–8748 (2022).

45. Kazak, L. et al. Alternative translation initiation augments the human mitochondrial proteome. Nucleic Acids Res 41, 2354–69 (2013).

46. Baker, A. R. & Slack, F. J. ADAR1 and its implications in cancer development and treatment. Trends Genet. 38, 821–830 (2022).

47. Sun, T. et al. Decoupling expression and editing preferences of ADAR1 p150 and p110 isoforms. Proc. Natl. Acad. Sci. 118, e2021757118 (2021).

48. Tsang, M.-J. & Cheeseman, I. M. Alternative CDC20 translational isoforms tune mitotic arrest duration. Nature 617, 154–161 (2023).

49. Higdon, A. L., Won, N. H. & Brar, G. A. Truncated protein isoforms generate diversity of protein localization and function in yeast. 2023.07.13.548938 Preprint at 10.1101/2023.07.13.548938 (2023).

50. Eisenberg, A. R. et al. Translation Initiation Site Profiling Reveals Widespread Synthesis of Non-AUG-Initiated Protein Isoforms in Yeast. Cell Syst. 11, 145–160.e5 (2020).

51. Sendoel, A. et al. Translation from unconventional 5′ start sites drives tumour initiation. Nature 541, 494–499 (2017).

52. Weber, R. et al. Monitoring the 5′UTR landscape reveals isoform switches to drive translational efficiencies in cancer. Oncogene 42, 638–650 (2023).

53. Starck, S. R. et al. Translation from the 5′ untranslated region shapes the integrated stress response. Science 351, aad3867 (2016).

54. Jackson, R. et al. The translation of non-canonical open reading frames controls mucosal immunity. Nature 564, 434–438 (2018).

55. Ivanov, I. P., Firth, A. E., Michel, A. M., Atkins, J. F. & Baranov, P. V. Identification of evolutionarily conserved non-AUG-initiated N-terminal extensions in human coding sequences. Nucleic Acids Res. 39, 4220–4234 (2011).

56. Damme, P. V., Gawron, D., Criekinge, W. V. & Menschaert, G. N-terminal Proteomics and Ribosome Profiling Provide a Comprehensive View of the Alternative Translation Initiation Landscape in Mice and Men *. Mol. Cell. Proteomics 13, 1245–1261 (2014).

57. Fournier, C. T. et al. Amino Termini of Many Yeast Proteins Map to Downstream Start Codons. J. Proteome Res. 11, 5712–5719 (2012).

58. Sherman, F. Getting started with yeast. in Methods in Enzymology (eds. Guthrie, C. & Fink, G. R.) vol. 350 3–41 (Academic Press, 2002).

59. Janke, C. et al. A versatile toolbox for PCR-based tagging of yeast genes: new fluorescent proteins, more markers and promoter substitution cassettes. Yeast 21, 947– 962 (2004).

60. Longtine, M. S. et al. Additional modules for versatile and economical PCR-based gene deletion and modification in Saccharomyces cerevisiae. Yeast 14, 953–961 (1998).

61. Storici, F. & Resnick, M. A. The Delitto Perfetto Approach to In Vivo Site-Directed Mutagenesis and Chromosome Rearrangements with Synthetic Oligonucleotides in Yeast. in Methods in Enzymology vol. 409 329–345 (Elsevier, 2006).

62. Chee, M. K. & Haase, S. B. New and Redesigned pRS Plasmid Shuttle Vectors for Genetic Manipulation of Saccharomyces cerevisiae. G3 GenesGenomesGenetics 2, 515–526 (2012).

63. Alberti, S., Gitler, A. D. & Lindquist, S. A suite of Gateway® cloning vectors for high-throughput genetic analysis in Saccharomyces cerevisiae. Yeast 24, 913–919 (2007).

64. Boulé, J.-B., Vega, L. R. & Zakian, V. A. The yeast Pif1p helicase removes telomerase from telomeric DNA. Nature 438, 57–61 (2005).

65. Wilson, M. A. et al. Pif1 helicase and Poldelta promote recombination-coupled DNA synthesis via bubble migration. Nature 502, 393–396 (2013).

66. Carreira, R., Aguado, F. J., Lama-Diaz, T. & Blanco, M. G. Holliday Junction Resolution. in Homologous Recombination: Methods and Protocols (eds. Aguilera, A. & Carreira, A.) 169–185 (Springer US, 2021). doi:10.1007/978-1-0716-0644-5_12.

67. Lee, B.-I. & Wilson, D. M. The RAD2 Domain of Human Exonuclease 1 Exhibits 5′ to 3′ Exonuclease and Flap Structure-specific Endonuclease Activities. J. Biol. Chem. 274, 37763–37769 (1999).

68. Sausen, C. W., Rogers, C. M. & Bochman, M. L. Thin-Layer Chromatography and Real-Time Coupled Assays to Measure ATP Hydrolysis. in DNA Repair (eds. Balakrishnan, L. & Stewart, J. A.) vol. 1999 245–253 (Springer New York, 2019).

69. Foster, S. S., Zubko, M. K., Guillard, S. & Lydall, D. MRX protects telomeric DNA at uncapped telomeres of budding yeast cdc13-1 mutants. DNA Repair 5, 840–851 (2006).

70. Eckert-Boulet, N., Rothstein, R. & Lisby, M. Cell Biology of Homologous Recombination in Yeast. in DNA Recombination (ed. Tsubouchi, H.) vol. 745 523–536 (Humana Press, 2011).

71. Kosugi, S., Hasebe, M., Tomita, M. & Yanagawa, H. Systematic identification of cell cycle-dependent yeast nucleocytoplasmic shuttling proteins by prediction of composite motifs. Proc. Natl. Acad. Sci. U. S. A. 106, 10171–10176 (2009).

72. Brameier, M., Krings, A. & MacCallum, R. M. NucPred—Predicting nuclear localization of proteins. Bioinformatics 23, 1159–1160 (2007).

73. Nguyen Ba, A. N., Pogoutse, A., Provart, N. & Moses, A. M. NLStradamus: a simple Hidden Markov Model for nuclear localization signal prediction. BMC Bioinformatics 10, 202 (2009).

74. Fukasawa, Y. et al. MitoFates: Improved Prediction of Mitochondrial Targeting Sequences and Their Cleavage Sites *[S]. Mol. Cell. Proteomics 14, 1113–1126 (2015).

75. Buchan, D. W. A. & Jones, D. T. The PSIPRED Protein Analysis Workbench: 20 years on. Nucleic Acids Res. 47, W402–W407 (2019).

