## Supplementary material for "Alternative translation initiation by ribosomal leaky scanning produces multiple isoforms of the Pif1 helicase": Supp Figures S1-S8 and Supp Tables S1-S3

Supp Fig 1

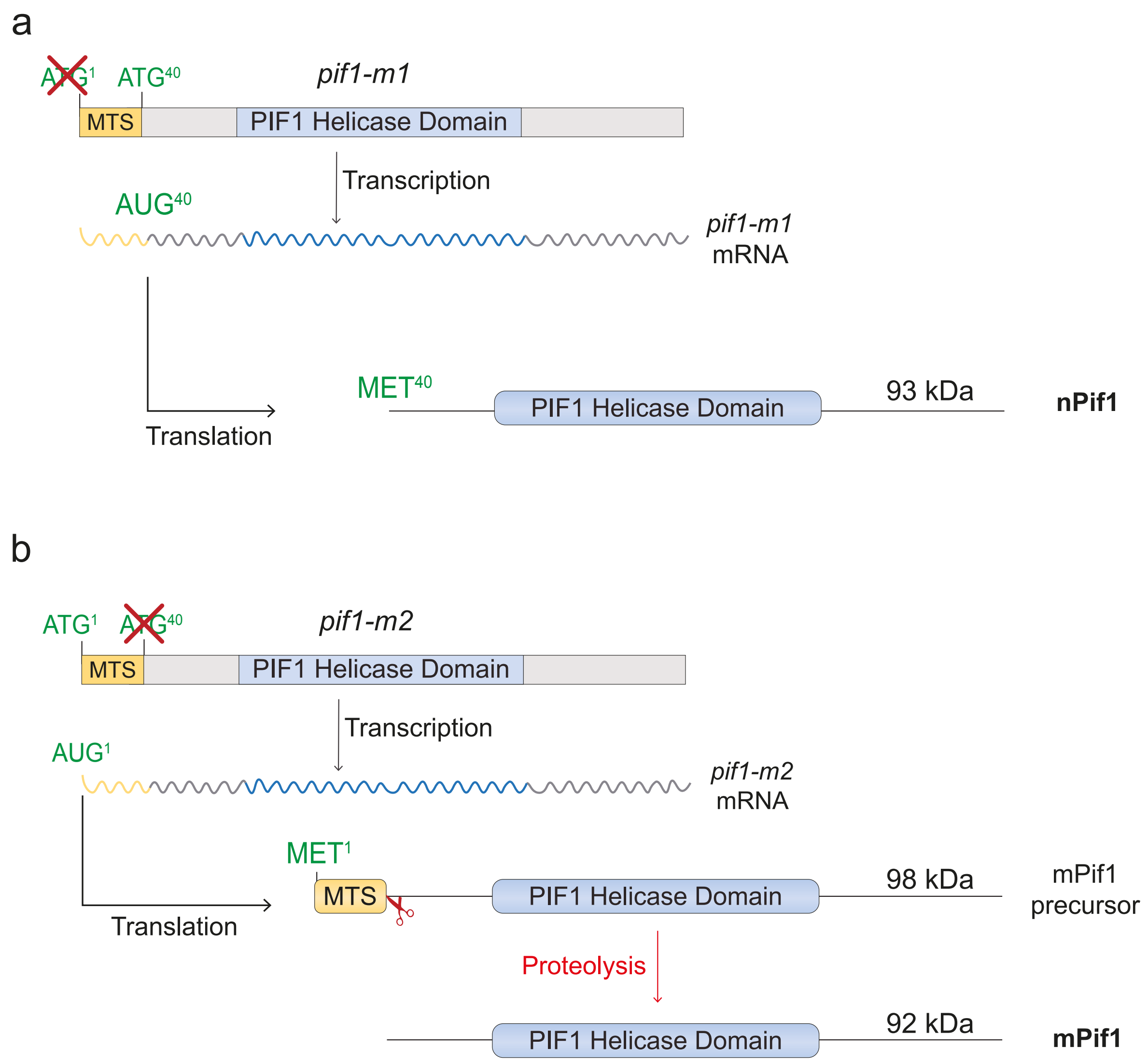


Supplementary Figure 1. **Classical separation-of-function alleles for nuclear and mitochondrial functions of *PIF1.***

(a) Schematic representation of *pif1-m1*, predicting the exclusive translation of the 93 kDa nuclear isoform nPif1.

(b) Schematic representation of *pif1-m2*, predicting the exclusive translation of the 98 kDa Pif1 mitochondrial precursor and its proteolytic processing into the mature, 92 kDa mitochondrial mPif1.

### Supp Fig 2

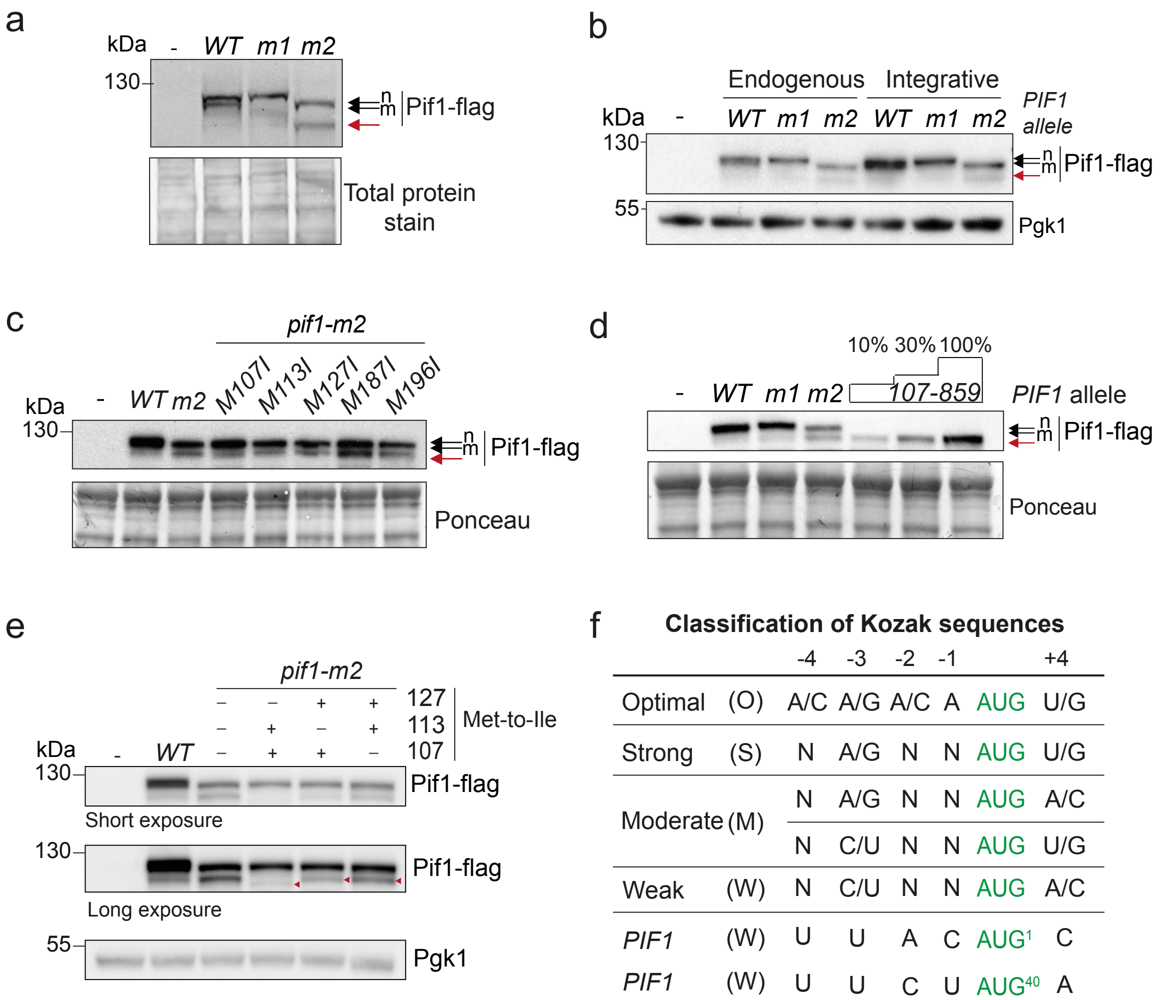


f **Classification of Kozak sequences**

| | | -4 | -3 | -2 | -1 | | +4 |
|---|---|---|---|---|---|---|---|
| Optimal | (O) | A/C | A/G | A/C | A | AUG | U/G |
| Strong | (S) | N | A/G | N | N | AUG | U/G |
| Moderate | (M) | N | A/G | N | N | AUG | A/C |
| | | N | C/U | N | N | AUG | U/G |
| Weak | (W) | N | C/U | N | N | AUG | A/C |
| *PIF1* | (W) | U | U | A | C | AUG[1] | C |
| *PIF1* | (W) | U | U | C | U | AUG[40] | A |

Supplementary Figure 2. **Characterization of the fast-migrating Pif1 isoform**.

(a) Assessment of the abundance of the fast-migrating Pif1 isoform in different *PIF1* mutants. Representative anti-FLAG fluorescent western blot of protein extracts from strains expressing untagged *PIF1* (-), *PIF1-6flag* (*WT*), *pif1-m1-6flag* (*m1*) or *pif1-m2-6flag* (*m2*), employed for the quantifications shown in Figure 2c. Black arrows indicate the nuclear (n) and mitochondrial Pif1 isoforms (m). The red arrow marks the fast-migrating isoform. Total protein stain was employed as a loading control (see Methods).

(b) Both strains expressing *PIF1* alleles from its endogenous locus or from an ectopic copy integrated at *ura3-52* exhibit the fast-migrating isoform. Anti-FLAG western blots of protein extracts from the indicated strains were carried out as in Figure 2e. Pgk1 was employed as a loading control. Arrows and labels as in (a).

(c) Single substitution of various Met codons to Ile codons at the indicated positions does not abrogate the production of the fast-migrating isoform. Western blot analysis of the indicated strains was carried out as in (b). Ponceau staining of the membrane was employed as a loading control. Arrows as in (a).

(d) An N-terminally truncated version of Pif1 (Pif1$^{107-859}$) co-migrates with the fast-migrating isoform enriched in *pif1-m2* mutants. Western blot analysis of the indicated strains was carried out as in (b). To carefully determine the migration of the Pif1$^{107-859}$ truncation with respect to the fast-migrating isoform, the protein extract of the *pif1*$^{107-859}$ strain was loaded undiluted (100%) or diluted (10% or 30%) in normalized extract from the untagged strain, thus avoiding distortion of migration due to salt effects. Ponceau staining of the membrane was employed as a loading control. Arrows as in (a).

(e) Double substitutions of Met to Ile codons indicate that Met$^{107}$ is the main translation initiation codon for the fast-migrating isoform. Western blot analysis of Pif1 in wild-type of *pif1-m2* strains harbouring the indicated single Met-to-Ile substitutions was performed as in (b). Arrowheads highlight the slightly different migration of the fast-migrating isoforms in different strains. Pgk1 was employed as a loading control.

(f) Classification of Kozak sequences in *S. cerevisiae* according to Meijer and Thomas[1]. The Kozak context for AUG$^{1}$ and AUG$^{40}$ in *PIF1* mRNA is shown below.

Supp Fig 3

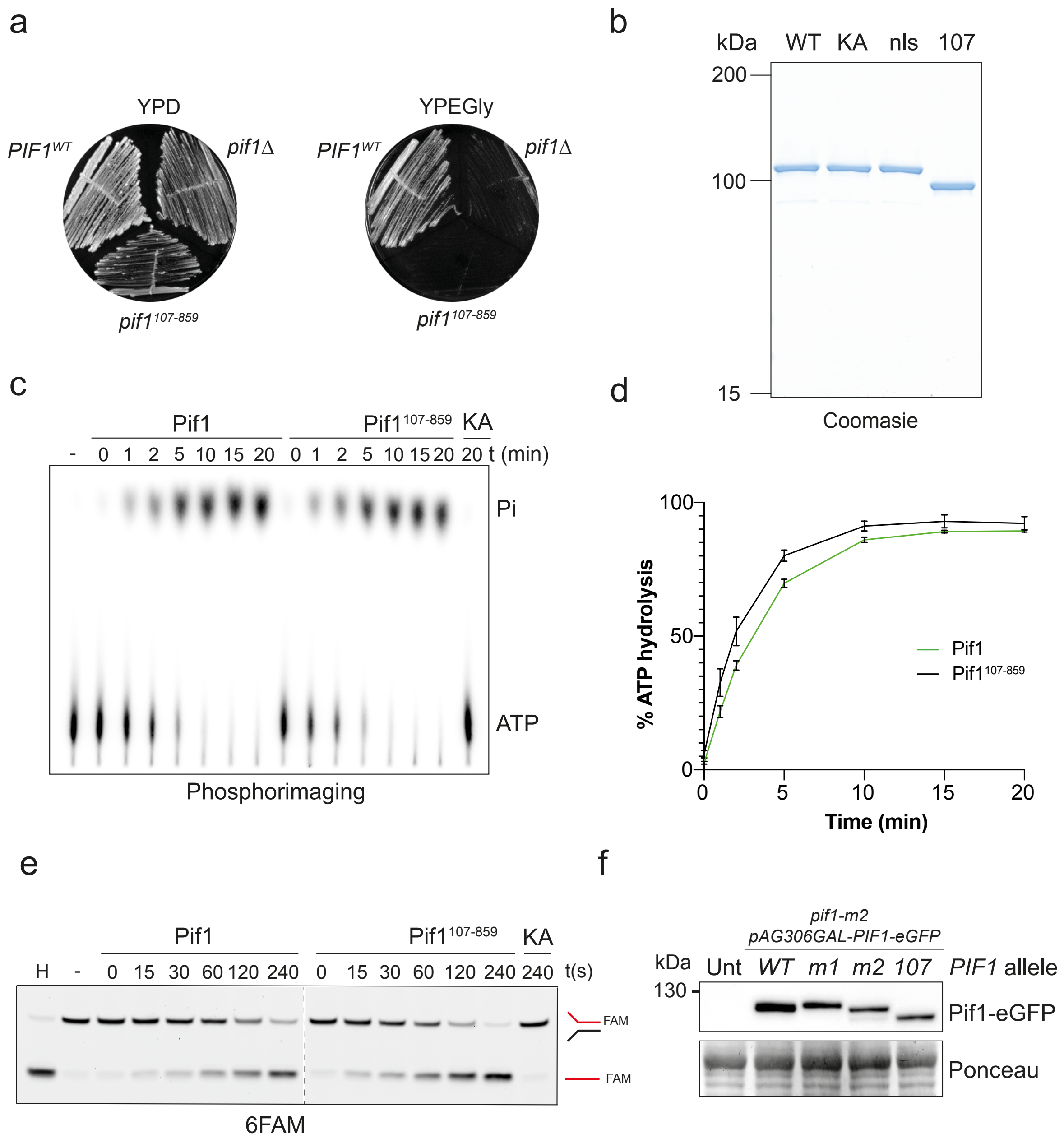
a
YPD
YPEGly
PIF1WT
pif1Δ
pif1107-859
b
kDa
WT
KA
nls
107
200
100
15
Coomasie
c
Pif1
Pif1107-859
KA
- 0 1 2 5 10 15 20 0 1 2 5 10 15 20 20 t (min)
Pi
ATP
Phosphorimaging
d
% ATP hydrolysis
100
50
0
0
5
10
15
20
Time (min)
Pif1
Pif1107-859
e
Pif1
Pif1107-859
KA
H - 0 15 30 60 120 240 0 15 30 60 120 240 240 t(s)
FAM
FAM
6FAM
f
pif1-m2
pAG306GAL-PIF1-eGFP
kDa
Unt
WT
m1
m2
107
PIF1 allele
130 -
Pif1-eGFP
Ponceau

Supplementary Figure 3. The N-terminally truncated Pif1$^{107\text{-}859}$ is an ATPase- and helicase-proficient Pif1 isoform.

(a) Pif1$^{107\text{-}859}$ cannot sustain growth on non-fermentable carbon sources. Strains harbouring the indicated *PIF1* alleles were streaked on rich medium with fermentable (YPD, 2% glucose) or non-fermentable (YPEGly, 3% ethanol, 3% glycerol) carbon sources. Plates were incubated at 30 ºC for 3 days and photographed.

(b) Purification of various Pif1 variants employed in this work. 1 µg of recombinant C-terminal 6xHis-tagged Pif1 (WT), Pif1$^{K264A}$ (KA), Pif1$^{nls}$ (nls) and Pif1$^{107\text{-}859}$ (107) was analysed by SDS-PAGE followed by Coomassie staining.

(c) Pif1$^{107\text{-}859}$ retains ATPase activity. ATP hydrolysis activity of recombinant Pif1 and Pif1$^{107\text{-}859}$ was determined by thin-layer chromatography using [$^{32}$P-$\gamma$]-ATP, followed by phosphorimaging. Catalytically inactive Pif1$^{KA}$ was employed as a negative control.

(d) Pif1$^{107\text{-}859}$ displays similar ATPase activity to Pif1. Quantification of ATP hydrolysis by Pif1 and Pif1$^{107\text{-}859}$ was estimated as percentage of inorganic phosphate release in time-course assays as in (c). Data are represented as mean values ± standard deviation from three replicates.

(e) Pif1$^{107\text{-}859}$ retains helicase activity. A representative DNA unwinding assay using 3'-6FAM-labelled splayed as a substrate and wild-type Pif1, Pif1$^{107\text{-}859}$ or Pif1$^{KA}$ (as a negative control) is shown. H, heat-denatured substrate.

(f) Overexpression of Pif1$^{107\text{-}859}$-eGFP from the inducible *GAL1* promoter is comparable to that of other Pif1 isoforms. After galactose induction, protein extracts from *pif1-m2* strains carrying an additional copy of a *PIF1* allele under the control of the *GAL1* promoter were analysed by Western blot using an anti-GFP antibody. Ponceau staining of the membrane was employed as a loading control.

### Supp Fig 4

a

| ***PIF1* alleles** | **Deleted region** |
|---|---|
| *Δ775-803* | EADEQVKKRKLDYAPGPKYKAKSKSKSNS |
| *Δ623-646* | KRVKTDDEVVLENIKRKEQLMQTI |
| *Δ650-662* | SAGKRRLPLVRFK |
| *Δ814-834* | NNGIAAMLQRHSRKRFQLKKE |

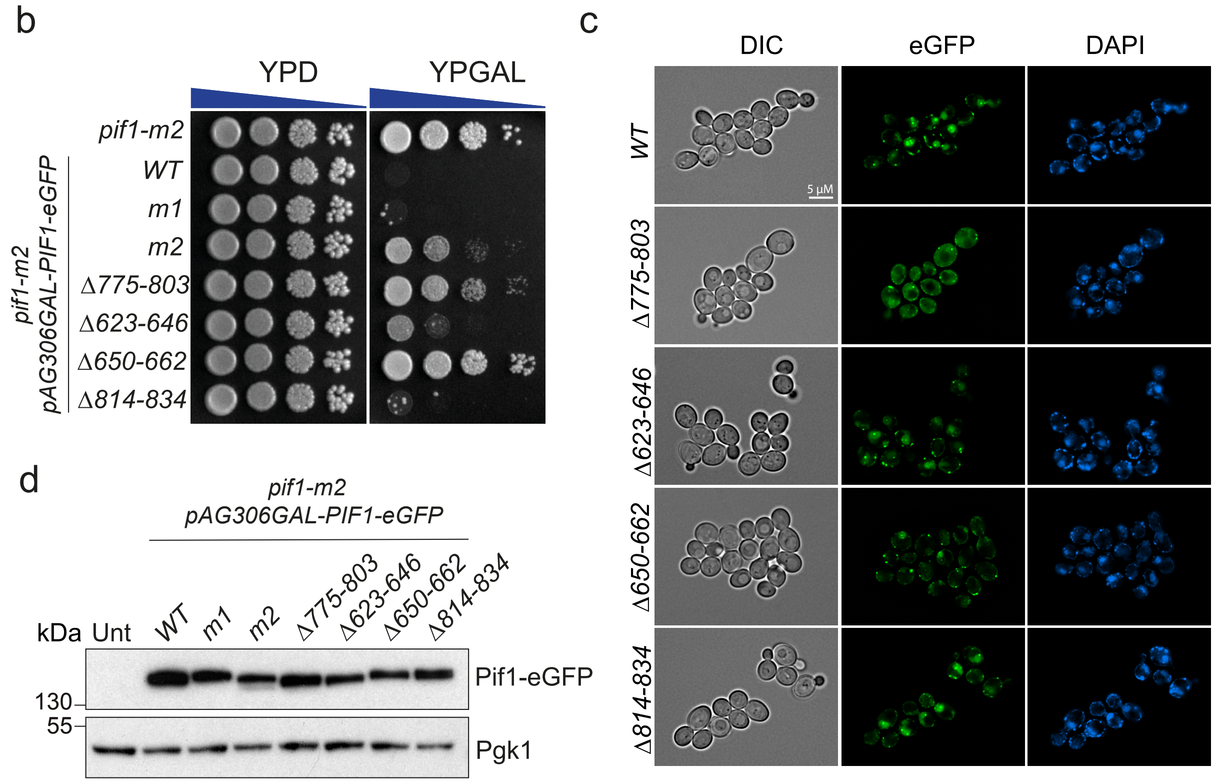

Supplementary Figure 4. **Characterization of potential NLS-containing regions in Pif1.**

(a) List of *PIF1* alleles with deletions in regions containing potential NLS sequences. Basic residues potentially relevant for nuclear import are indicated in green.

(b) Deletion of the 775-803 and 650-659 regions counteracts the lethality induced by Pif1 overexpression. Ten-fold dilutions of strains with the indicated genotypes were plated on YPD or YPGal plates and imaged after 2 days of incubation at 30 ºC.

(c) Deletion of the 775-803 region in Pif1 impairs its nuclear import. Subcellular localization of Pif1 isoforms in strains overexpressing different *PIF1-eGFP* alleles was assessed in live cells by fluorescence microscopy. DNA was visualized with DAPI staining and cell contour by DIC microscopy.

(d) Expression levels of overexpressed Pif1 deletion mutants are comparable. Abundance of Pif1 variants in the strains from (b) and (c) was estimated by western blotting using anti-GFP antibodies. Protein extracts were prepared from liquid cultures grown in YPRaff and induced for 3 h by addition of 2% galactose. Pgk1 was used as a loading control.

### Supp Fig 5

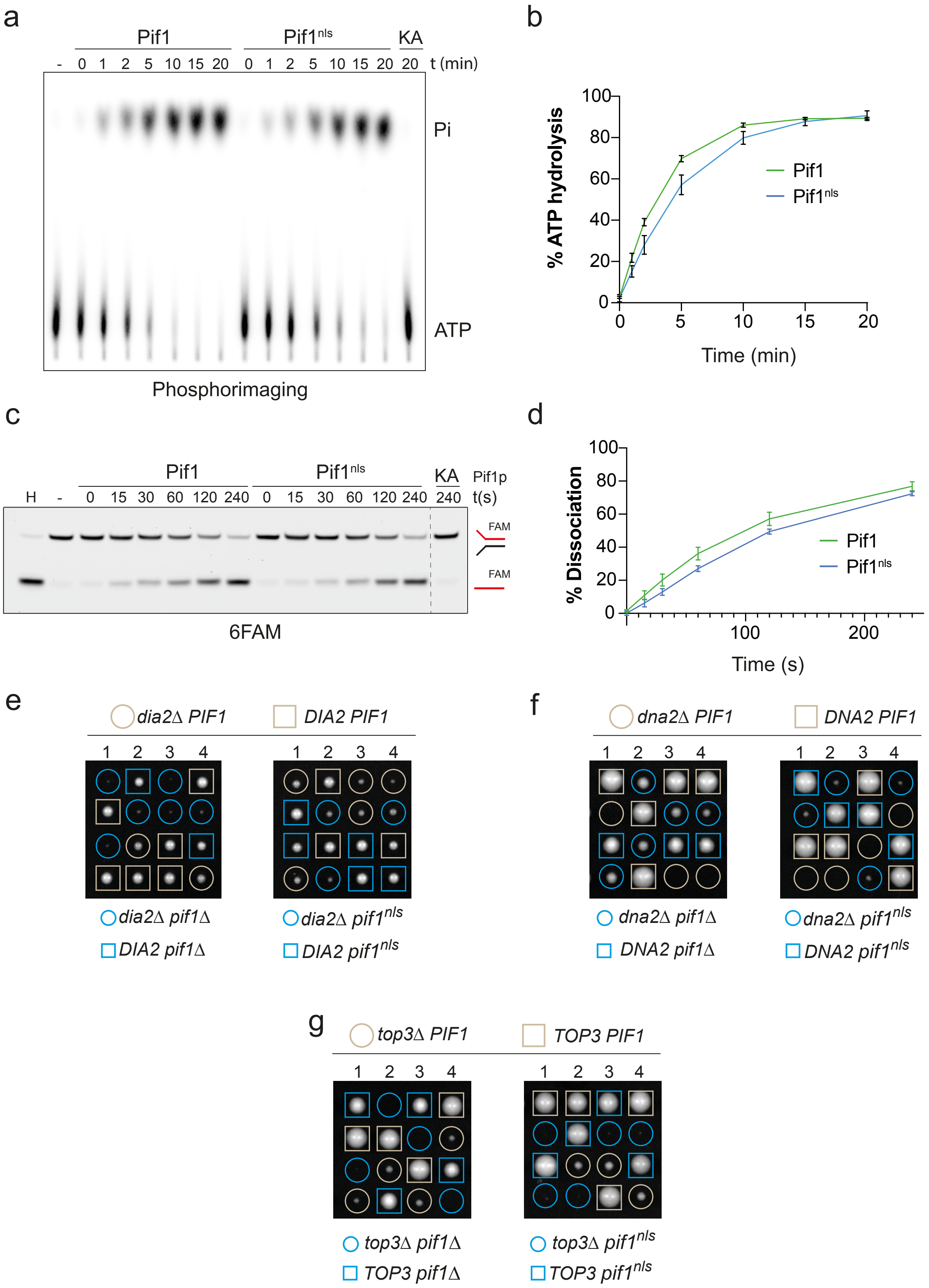
a
Pif1
Pif1nls
KA
- 0 1 2 5 10 15 20 0 1 2 5 10 15 20 20 t (min)
Pi
ATP
Phosphorimaging
b
% ATP hydrolysis
Time (min)
Pif1
Pif1nls
c
Pif1
Pif1nls
KA
Pif1p
H - 0 15 30 60 120 240 0 15 30 60 120 240 240 t(s)
FAM
FAM
6FAM
d
% Dissociation
Time (s)
Pif1
Pif1nls
e
dia2Δ PIF1
DIA2 PIF1
1 2 3 4
dia2Δ pif1Δ
DIA2 pif1Δ
dia2Δ pif1nls
DIA2 pif1nls
f
dna2Δ PIF1
DNA2 PIF1
1 2 3 4
dna2Δ pif1Δ
DNA2 pif1Δ
dna2Δ pif1nls
DNA2 pif1nls
g
top3Δ PIF1
TOP3 PIF1
1 2 3 4
top3Δ pif1Δ
TOP3 pif1Δ
top3Δ pif1nls
TOP3 pif1nls

Supplementary Figure 5. ***pif1**$^{nls}$* **strains retain phenotypically relevant amounts of nuclear Pif1 activity.**

(a) Pif1$^{nls}$ retains ATPase activity. ATP hydrolysis activity of recombinant Pif1 and Pif1$^{107\text{-}859}$ was determined by thin-layer chromatography using [$^{32}$P-$\gamma$]-ATP, followed by phosphorimaging. Catalytically inactive Pif1$^{KA}$ was employed as a negative control.

(b) Pif1$^{nls}$ displays similar ATPase activity to Pif1. Quantification of ATP hydrolysis by Pif1 and Pif1$^{nls}$ was estimated as percentage of inorganic phosphate release in time-course assays as in (a). Data are represented as mean values ± standard deviation from three replicates.

(c) Pif1$^{nls}$ retains helicase activity. A representative DNA unwinding assay using a 3'-6FAM-labelled splayed arm as a substrate and wild-type Pif1, Pif1$^{nls}$ or Pif1$^{KA}$ (as a negative control) is shown. H, heat-denatured substrate.

(d) Helicase activity of Pif1$^{nls}$ *in vitro* is comparable to wild-type Pif1. Quantification of ATP hydrolysis by Pif1 and Pif1$^{nls}$ was estimated as percentage of substrate dissociation in time-course assays from three replicates as in (c). Data are represented as mean values ± standard deviation from three replicates.

(e) Tetrad microdissection of diploid strains carrying heterozygous mutations for the indicated wild-type and mutant alleles of *DIA2* and *PIF1*. Images were taken after 2 days of incubation at 30 ºC.

(f) Tetrad microdissection of diploid strains carrying heterozygous mutations for the indicated wild-type and mutant alleles of *DNA2* and *PIF1*. Images were taken after 2 days of incubation at 30 ºC.

(g) Tetrad microdissection of diploid strains carrying heterozygous mutations for the indicated wild-type and mutant alleles of *TOP3* and *PIF1*. Images were taken after 3 days of incubation at 30 ºC.

Supp Fig 6

a

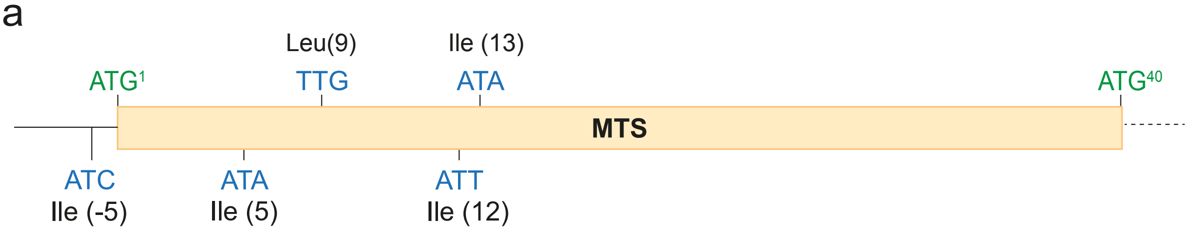


b

| Start codon | Probability of MTS |
|---|---|
| AUC-Ile (-5) | 0.844 |
| AUG-Met (1) | 0.996 |
| AUA-Ile (5) | 0.911 |
| UUG-Leu (9) | 0.627 |
| AUU-Ile (12) | 0.465 |
| AUA-Ile (13) | 0.383 |

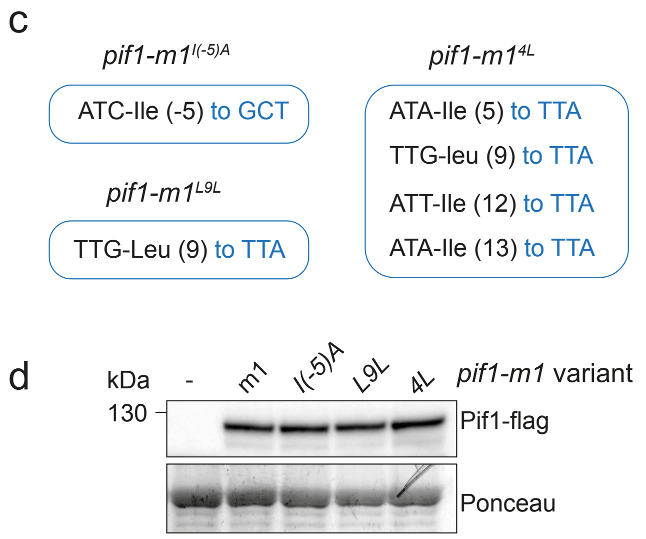


Supplementary Figure 6. **Identification of near-cognate start codons in *PIF1* with potential to generate Pif1 isoforms with a functional MTS.**

(a) Schematic representation of the near-cognate start codons upstream and downstream ATG[1] in *PIF1*.

(b) Probability of MTS in N-terminal truncated or elongated Pif1 isoforms translated from near-cognates depicted in (a). Scores obtained from Mitofates (see *Methods*).

(c) List of conservative substitutions introduced in near-cognate ATGs in the *pif1-m1* allele to generate the indicated alleles.

(d) Expression levels of *pif1-m1* variants with near-cognate substitutions are expressed at similar levels. Western blot analysis of Pif1-flag in protein extracts from strains ectopically expressing the indicated *pif1-m1* variants. (-), untagged *pif1-m1* strain. Ponceau staining of the PVDF membrane was employed as a loading control.

### Supp Fig 7

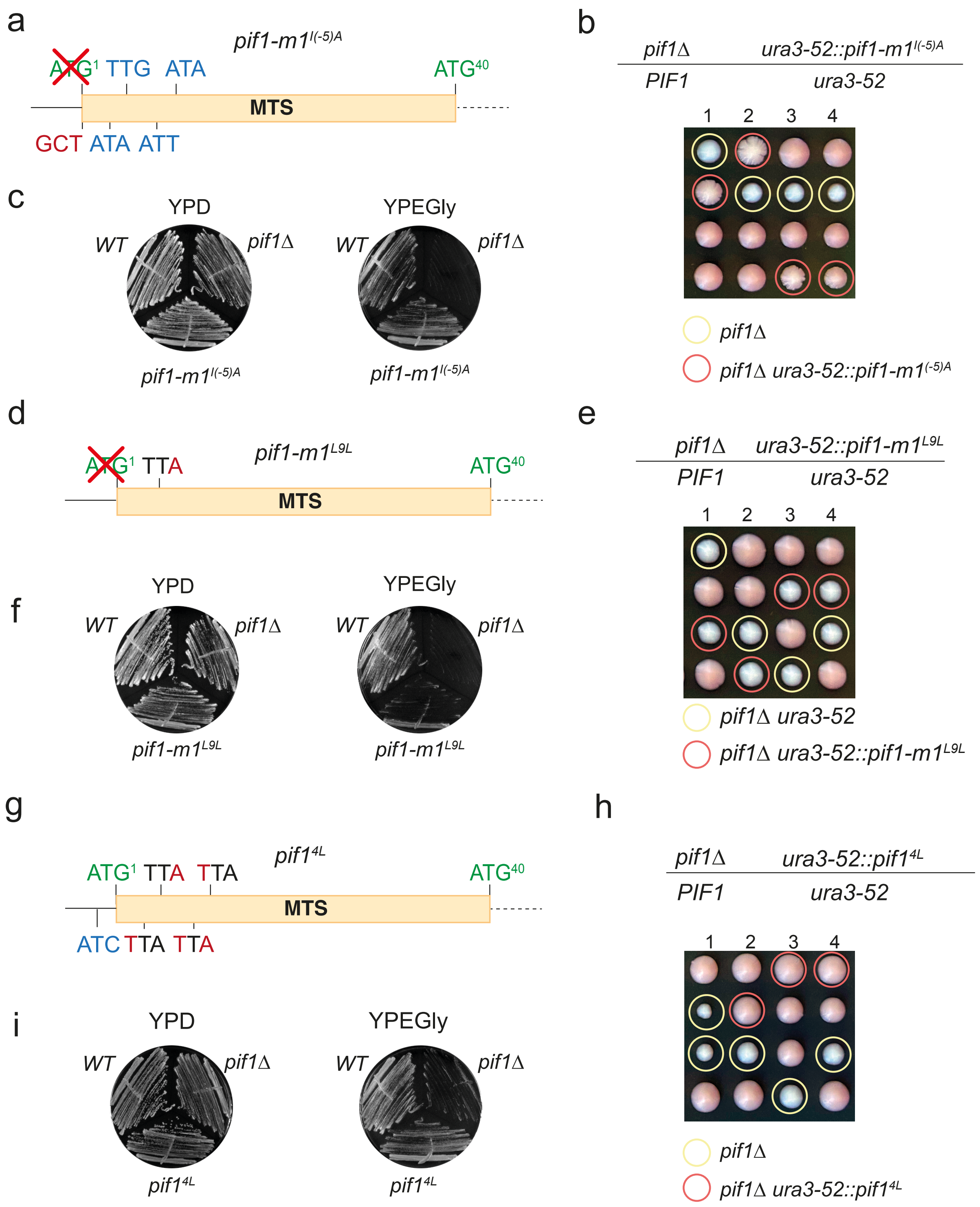
a
pif1-m1^I(-5)A
ATG¹ TTG ATA
ATG⁴⁰
MTS
GCT ATA ATT
b
pif1Δ ura3-52::pif1-m1^I(-5)A
PIF1 ura3-52
1 2 3 4
pif1Δ
pif1Δ ura3-52::pif1-m1^I(-5)A
c
YPD
YPEGly
WT
pif1Δ
pif1-m1^I(-5)A
d
pif1-m1^L9L
ATG¹ TTA
ATG⁴⁰
MTS
e
pif1Δ ura3-52::pif1-m1^L9L
PIF1 ura3-52
1 2 3 4
pif1Δ ura3-52
pif1Δ ura3-52::pif1-m1^L9L
f
YPD
YPEGly
WT
pif1Δ
pif1-m1^L9L
g
pif1^4L
ATG¹ TTA TTA
ATG⁴⁰
MTS
ATC TTA TTA
h
pif1Δ ura3-52::pif1^4L
PIF1 ura3-52
1 2 3 4
pif1Δ
pif1Δ ura3-52::pif1^4L
i
YPD
YPEGly
WT
pif1Δ
pif1^4L

Supplementary Figure 7. **Differential contribution by near-cognate start codons near ATG$^{1}$ for the production of functional mPif1**.

(a) Schematic representation of the N-terminal MTS region of the *pif1-m1*$^{I(-5)A}$, depicting the start codon (green), the mutations in the upstream near-cognate start codon Ile$^{-5}$ (red) and unmodified downstream near cognates (blue).

(b) *pif1-m1*$^{I(-5)A}$ strains do not display an overt *petite* phenotype after tetrad dissection. Tetrads from a diploid strain carrying a heterozygous deletion of *PIF1* and an ectopic *pif1-m1*$^{I(-5A)}$ allele integrated at *ura3-52* were microdissected on YPD plates and incubated at 30 ºC for 3 days. Adenine was omitted from YPD to allow for colour discrimination of *petite* colonies.

(c) Non-fermentable carbon sources can sustain growth of *pif1-m1*$^{I(-5)A}$ strains. Freshly generated strains harbouring the indicated *PIF1* alleles were streaked on rich medium with a fermentable (YPD, 2% glucose) or non-fermentable (YPEGly, 3% ethanol, 3% glycerol) carbon source. Plates were incubated at 30 ºC for 3 days and photographed.

(d) Schematic representation of the N-terminal MTS region of the *pif1-m1*$^{L9L}$, depicting the start codon (green), and mutation in the downstream near-cognate start codon Leu$^{9}$ (red).

(e) *pif1-m1*$^{L9L}$ strains display a clear *petite* phenotype after tetrad dissection. Tetrad dissections were carried out as in (b).

(f) Non-fermentable carbon sources sustain residual growth of *pif1-m1*$^{L9L}$ strains. Strains were tested as in (c).

(g) Schematic representation of the N-terminal MTS region of the *pif1*$^{4L}$, depicting the start codon (green), the upstream near-cognate start codon (blue) and mutations in the four downstream near-cognate start codons (red).

(h) *pif1*$^{4L}$ strains do not display a *petite* phenotype after tetrad dissection. Tetrad dissections were carried out as in (b).

(i) Non-fermentable carbon sources sustain growth of *pif1*$^{4L}$ strains. Strains were tested as in (c).

Supp Fig 8

a

| Near cognate | Position | -3 position |
|---|---|---|
| AUU | 51 | Pyrimidine |
| UUG | 55 | Pyrimidine |
| AUA | 60 | Pyrimidine |
| UUG | 68 | Purine |
| AUA | 81 | Pyrimidine |
| AUU | 92 | Purine |
| AUC | 93 | Purine |
| AAC | 96 | Purine |
| CUG | 102 | Pyrimidine |

b

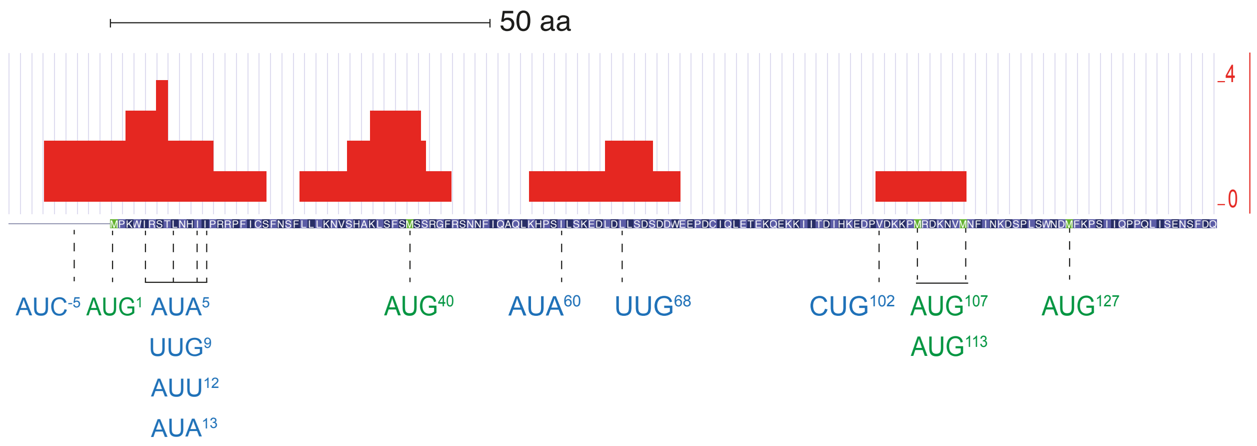


Supplementary Figure 8. **Near-cognate start codons downstream AUG$^{40}$ may constitute a source of additional nuclear Pif1 isoforms**.

(a) List of near-cognate start codons between positions 40-107 in *PIF1*. The presence of a purine in -3 position favours ATI from near-cognate start codons[2].

(b) Small ribosome scanning subunits can be detected at near-cognate codons that could produce alternative N-terminally truncated Pif1 isoforms. TCP-seq data from Archer et al.[3] was visualized employing GWIPS-viz[4]. The positions of relevant AUG (green) and near-cognate start codons (blue) is indicated. Y-axis represents the footprint counts of small ribosome subunits.

Supplementary Table S1. **Yeast strains employed in this work.**

| Number | Genotype |
|---|---|
| MGBY597 | *MATa pif1Δ::HIS3MX6* |
| MGBY665[5] | *MATa pif1-m2* |
| MGBY760 | *MATa PIF1-WT-6xHis-6xFlag::klTRP1* |
| MGBY764 | *MATa pif1-m1-6xHis-6xFlag::klTRP1* |
| MGBY768 | *MATa pif1-m2-6xHis-6xFlag::klTRP1* |
| MGBY834 | *MATa pif1-m2-6xHis-6xFlag::klTRP1 rad3-102::hyg* |
| MGBY835 | *MATa rad3-102::hyg* |
| MGBY841* | *MATa PIF1-6xHis-6xFlag::klTRP1* |
| MGBY844* | *MATa pif1-m1-6xHis-6xFlag::klTRP1* |
| MGBY847* | *MATa pif1-m2-6xHis-6xFlag::klTRP1* |
| MGBY886 | *MATa/MATalpha*<br>*Heterozygous for:*<br>*pif1-m2-6xHis-6xFlag::klTRP1 / PIF1*<br>*dia2Δ::kanMX / DNA2* |
| MGBY924 | *MATa pif1-m2 ura3-1::pRSII406-PIF1-WT-6xHis-6xFlag* |
| MGBY927 | *MATa pif1-m2 ura3-1::pRSII406-pif1-m1-6xHis-6xFlag* |
| MGBY930 | *MATa pif1-m2 ura3-1::pRSII406-pif1-m2-6xHis-6xFlag* |
| MGBY933 | *MATa pif1-m2 ura3-1::pRSII406-PIF1-WT-K1-6xHis-6xFlag* |
| MGBY936 | *MATa pif1-m2 ura3-1::pRSII406-PIF1-WT-K40-6xHis-6xFlag* |
| MGBY942 | *MATa pif1-m2 ura3-1::pRSII406-pif1-m1-K40-6xHis-6xFlag* |
| MGBY967 | *MATa pif1-m2 ura3-1::pRSII406-pif1-m2-M107I-6xHis-6xFlag* |
| MGBY970 | *MATa pif1-m2 ura3-1::pRSII406-pif1-m2-M113I-6xHis-6xFlag* |
| MGBY972 | *MATa pif1-m2 ura3-1::pRSII406-pif1-m2-M127I-6xHis-6xFlag* |
| MGBY975 | *MATa pif1-m2 ura3-1::pRSII406-pif1-m2-M187I-6xHis-6xFlag* |
| MGBY978 | *MATa pif1-m2 ura3-1::pRSII406-pif1-m2-M196I-6xHis-6xFlag* |
| MGBY1091 | *MATa/MATalpha*<br>*Heterozygous for:*<br>*kanMX::*$P_{ADH1}$*-pif1-m2-6xHis-6xFlag::klTRP1 / pif1-m2-6xHis-6xFlag::klTRP1*<br>*dna2Δ::LEU2 / DNA2* |
| MGBY1097 | *MATa pif1Δ::HIS3MX6* $P_{PIF1}$*::pif1-107-859-6xHis-6xFlag::pRSII406* |
| MGBY1108 | *MATa / MATalpha*<br>*Heterozygous for*<br>*pif1Δ::HIS3MX6* $P_{PIF1}$*::pif1-107-859-6xHis-6xFlag::pRSII406 / PIF1*<br>*dia2Δ::kanMX / DIA2* |
| MGBY1111 | *MATa pif1Δ::HIS3MX6* $P_{PIF1}$*::PIF1-107-6xHis-6xFlag::pRSII406* |
| MGBY1115 | *MATa rad3-102::hyg pif1Δ::HIS3MX6 PIF1-107-6xHis-6xFlag::pRSII406* |
| MGBY1162 | *MATa pif1-m2 ura3-1::pRSII406-pif1-m2-M107I-M113I-M127I-6xHis-6xFlag* |
| MGBY1178 | *MATa/MATalpha*<br>*Heterozygous for:*<br>*pif1Δ::HIS3MX6/PIF1*<br>*top3Δ::LEU2/TOP3* |

| | |
|---|---|
| MGBY1180 | MATa/MATalpha<br>Heterozygous for:<br>pif1-m1-6xHis-6xFlag::klTRP1/PIF1<br>top3Δ::LEU2/TOP3 |
| MGBY1182 | MATa/MATalpha<br>Heterozygous for:<br>pif1-m2-6xHis-6xFlag::klTRP1/PIF1<br>top3Δ::LEU2/TOP3 |
| MGBY1193 | MATa/MATalpha<br>Heterozygous for:<br>dia2Δ::kanMX/DIA2<br>pif1Δ::his3MX6/PIF1 |
| MGBY1194 | MATa/MATalpha<br>Heterozygous for:<br>dia2Δ::kanMX/DIA2<br>pif1-m1-6xHis-6xFlag::klTRP1/PIF1 |
| MGBY1203 | MATa pif1-m2 ura3-1::pAG306GAL-775-803-NLS-PIF1-eGFP |
| MGBY1211 | MATA pif1-m2 ura3-1::pAG306GAL-775-803-NLS-4A-eGFP |
| MGBY1214 | MATa pif1-m2 ura3-1::pAG306GAL-MET-linker-eGFP |
| MGBY1215 | MATa |
| MGBY1230 | MATa/MATalpha<br>Heterozygous for:<br>kanMX::$P_{ADH1}$-pif1-m2-6xHis-6xFlag::klTRP1 / pif1-m2-6xHis-6xFlag::klTRP1<br>top3Δ::LEU2/ TOP3 |
| MGBY1234 | MATa pif1-m2 ura3-1:pRSII406-pif1-107-859-6xHis-6xFlag |
| MGBY1237 | MATa/MATalpha<br>Heterozygous for:<br>pif1Δ::HIS3MX6 / PIF1<br>dna2Δ::LEU2 / DNA2 |
| MGBY1239 | MATa/MATalpha<br>Heterozygous for:<br>pif1-m1-6xHis-6xFlag::klTRP1 /PIF1<br>dna2Δ::LEU2 / DNA2 |
| MGBY1242 | MATa/MATalpha<br>Heterozygous for:<br>pif1-m2-6xHis-6xFlag::klTRP1 / PIF1<br>dna2Δ::LEU2 / DNA2 |
| MGBY1273 | MATa/MATalpha<br>Heterozygous for:<br>pif1Δ::HIS3MX6 pPIF1::pif1-107-859-6xHis-6xFlag::pRSII406 / PIF1<br>top3Δ::LEU2 / TOP3 |
| MGBY1274 | MATa/MATalpha<br>Heterozygous for:<br>pif1Δ::HIS3MX6 pPIF1::pif1-107-859-6xHis-6xFlag::pRSII406 / PIF1<br>dna2Δ::LEU2 / DNA2 |
| MGBY1278 | MATa<br>Heterozygous for:<br>pif1-nls-6xHis-6xFlag::klTRP1 / PIF1<br>top3Δ::LEU2 / TOP3 |
| MGBY1280 | MATa/MATalpha<br>Heterozygous for: |

| | |
|---|---|
| | *pif1-nls-6xHis-6xFlag::klTRP1 / PIF1*<br>*dna2∆::LEU2 / DNA2* |
| MGBY1283 | *MATa/MATalpha*<br>*Heterozygous for:*<br>*pif1-nls-6xHis-6xFlag::klTRP1 / PIF1*<br>*dia2∆::kanMX / DIA2* |
| MGBY1319 | *MATa/MATalpha*<br>*Heterozygous for:*<br>*pif1∆::HIS3MX6 / PIF1*<br>*ura3-52::pRSII406-pif1-m1-6xHis-6xFlag / ura3-52* |
| MGBY1321 | *MATa/MATalpha*<br>*Heterozygous for:*<br>*pif1∆::HIS3MX6 / PIF1*<br>*ura3-52::pRSII406-pif1-m1-IL(-5)A-6xHis-6xFlag / ura3-52* |
| MGBY1323 | *MATa/MATalpha*<br>*Heterozygous for:*<br>*pif1∆::HIS3MX6 / PIF1*<br>*ura3-52::pRSII406-pif1-m1-L9L-6xHis-6xFlag / ura3-52* |
| MGBY1326 | *MATa/MATalpha*<br>*Heterozygous for:*<br>*pif1∆::HIS3MX6 / PIF1*<br>*ura3-52::pRSII406-pif1-m1-4L-6xHis-6xFlag / ura3-52* |
| MGBY1352 | *MATa pif1-mit-6xHis-6xFlag::klTRP1* |
| MGBY1356 | *MATa pif1-m1-L9L-6xHis-6xFlag::klTRP1* |
| MGBY1374 | *MATa/MATalpha*<br>*Heterozygous:*<br>*pif1∆::LEU2*<br>*ura3-52::pRSII406-PIF1-WT-6xHis-6xFlag* |
| MGBY1377 | *MATa/MATalpha*<br>*Heterozygous:*<br>*pif1∆::LEU2 / PIF1*<br>*ura3-52::pRSII406-pif1-m1-6xHis-6xFlag / ura3-52* |
| MGBY1378 | *MATa/MATalpha*<br>*Heterozygous:*<br>*pif1∆::LEU2 / PIF1*<br>*ura3-52::pRSII406-pif1-m2-6xHis-6xFlag / ura3-52* |
| MGBY1383 | *MATa/MATalpha*<br>*Heterozygous:*<br>*pif1∆::LEU2*<br>*ura3-52::pRSII406-pif1-nls-6xHis-6xFlag / ura3-52* |
| MGBY1386 | *MATa/MATalpha*<br>*Heterozygous:*<br>*pif1∆::LEU2*<br>*ura3-52::pRSII406-pif1-mit-6xHis-6xFlag / ura3-52* |
| MGBY1415 | *MATa/MATalpha*<br>*Heterozygous:*<br>*pif1-mit-6xHis-6xFlag::klTRP1*<br>*dna2∆::LEU2* |
| MGBY1421 | *MATa rad3-102::hyg* |
| MGBY1426 | *MATalpha rad3-102::hyg pif1-mit-6xHis-6xFlag::klTRP1* |
| MGBY1492 | *MATa/MATalpha*<br>*Heterozygous for:* |

| | |
|---|---|
| | *dia2Δ::kanMX / DIA2*<br>*pif1Δ::his3MX / pif1-mit-6xHis-6xFlag::klTRP1* |
| MGBY1496 | *MATa pif1Δ::HIS3MX6 ura3-52::pRSII406-pif1-4L-6xHis-6xFlag* |
| MGBY1527 | *MATa rad3-102::hyg pif1Δ::HIS3MX6* |
| MGBY1536 | *MATa rad3-102::hyg pif1-m1-6xHis-6xFlag::klTRP1* |
| MGBY1579 | *MATa/MATalpha*<br>*Heterozygous for:*<br>*dia2Δ::kanMX /DIA2*<br>*pif1-m2-6xHis-6xFlag::klTRP1 / pif1-mit-6xHis-6xFlag::klTRP1* |
| MGBY1581 | *MATa/MATalpha*<br>*Heterozygous for:*<br>*dna2Δ::LEU2 / DNA2*<br>*pif1-m2-6xHis-6xFlag::klTRP1 / pif1-mit-6xHis-6xFlag::klTRP1* |
| MGBY1702 | *pif1-m2 ura3-1::pAG306GAL-PIF1-eGFP* |
| MGBY1705 | *pif1-m2 ura3-1::pAG306GAL-pif1-m1-eGFP* |
| MGBY1706 | *pif1-m2 ura3-1::pAG306GAL-pif1-m2-eGFP* |
| MGBY1709 | *pif1-m2 ura3-1::pAG306GAL-pif1-Δ775-803-eGFP* |
| MGBY1713 | *pif1-m2 ura3-1::pAG306GAL-pif1-Δ623-646-eGFP* |
| MGBY1713 | *pif1-m2 ura3-1::pAG306GAL-pif1-Δ650-662-eGFP* |
| MGBY1714 | *pif1-m2 ura3-1::pAG306GAL-pif1-Δ814-834-eGFP* |
| MGBY1717 | *pif1-m2 ura3-1::pAG306GAL-pif1-9A-eGFP* |
| MGBY1718 | *pif1-m2 ura3-1::pAG306GAL-pif1-4A-eGFP* |
| MGBY1721 | *pif1-m2 ura3-1::pAG306GAL-pif1-5A-eGFP* |
| MGBY1722 | *pif1-m2 ura3-1::pAG306GAL-pif1-107-859-eGFP* |
| MGBY1724 | *MATa pif1-m2 ura3-1::pRSII406-pif1-m2-M107I-M113I-6xHis-6xFlag* |
| MGBY1726 | *MATa pif1-m2 ura3-1::pRSII406-pif1-m2-M113I-M127I-6xHis-6xFlag* |
| MGBY1729 | *MATa pif1-m2 ura3-1::pRSII406-pif1-m2-M107I-M127II-6xHis-6xFlag* |

All strains are derivatives of W303 background (*ura3-52; trp1Δ2; leu2-3,112; his3-11; ade2-1; can1-100; RAD5*+) or *(ura3-1; trp1-1, leu2-3,112; his3-11,15; ade2-1; can1-100; RAD5+).*

*YPH background (*ura3-52; lys2-801_amber; ade2-101_ochre; trp1-Δ63; his3-Δ200; leu2-Δ1*).

Supplementary Table S2**. Plasmids employed in this study.**

| **Alias** | **Name** | **Source** |
|---|---|---|
| pMB1 | pUC19-6xHis-6xFlag::klTRP1 | Kind gift of Prof. Joao Matos |
| pMB34 | pFA6a-LEU2 | Houseley and Tollervey[7] |
| pMB40 | pFA6a-kanMX4 | Watch et al.[8] |
| pMB41 | pFA6a-HIS3MX6 | Longtine et al[9] |
| pMB42 | pCORE-UH | Storici and Resnick[10] |
| pMB198 | pAG306GAL-ccdB-eGFP | Alberti et al.[11] |
| pMB206 | pRSII406-PIF1-6xHis-6xFlag | This work |
| pMB207 | pRSII406-pif1-m1-6xHis-6xFlag | This work |
| pMB208 | pRSII406-pif1-m2-6xHis-6xFlag | This work |
| pMB217 | pRSII406-pif1-m2-M107I-6xHis-6xFlag | This work |
| pMB218 | pRSII406-pif1-m2-M113I-6xHis-6xFlag | This work |
| pMB219 | pRSII406-pif1-m2-M127I-6xHis-6xFlag | This work |
| pMB220 | pRSII406-pif1-m2-M187I-6xHis-6xFlag | This work |
| pMB221 | pRSII406-pif1-m2-M196I-6xHis-6xFlag | This work |
| pMB223 | pRSII406-pif1-Kozak-MET40-6xHis-6xFlag | This work |
| pMB225 | pRSII406-pif1-M1-Kozak-MET40-6xHis-6xFlag | This work |
| pMB236 | pET28b-6xHis-nPIF1 | Boulé et al.[12] Addgene (#65047) |
| pMB237 | pET28b-6xHis-nPIF1-K264A | This work |
| pMB238 | pET28b-6xHis-nPIF1-nls | This work |
| pMB242 | pRSII406-pif1-m2-M107I-M127I-6xHis-6xFlag | This work |
| pMB246 | pRSII406-pif1-M107I-M113I-M127I-6xHis-6xFlag | This work |
| pMB247 | pET28b-6xHis-pif1-107-859 | This work |
| pMB248 | pRSII406-pif1-107-859-6xHis-6xFlag | This work |
| pMB250 | pBluescriptII-yeast-telomere | Kind gift of David Lydall |
| pMB274 | pAG306GAL-PIF1-eGFP | This work |
| pMB275 | pAG306GAL-pif1-m1-eGFP | This work |
| pMB276 | pAG306GAL-pif1-m2-eGFP | This work |
| pMB277 | pAG306GAL-pif1-Δ775-803-eGFP | This work |
| pMB278 | pAG306GAL-pif1-4A-eGFP | This work |
| pMB279 | pAG306GAL-pif1-5A-eGFP | This work |

| | | |
|---|---|---|
| pMB281 | pAG306GAL-pif1-9A-eGFP | This work |
| pMB284 | pAG306GAL-pif1-Δ623-646-eGFP | This work |
| pMB286 | pAG306GAL-pif1-Δ650-662-eGFP | This work |
| pMB288 | pAG306GAL-pif1-Δ814-834-eGFP | This work |
| pMB290 | pAG306GAL-pif1-107-859-eGFP | This work |
| pMB303 | pRSII406-pif1-M107I-M113I-M127I-6xHis-6xFlag | This work |
| pMB306 | pAG306GAL-775-803-NLS-PIF1-eGFP | This work |
| pMB313 | pAG306GAL-775-803-NLS-4A-PIF1-eGFP | This work |
| pMB315 | pAG306GAL-MET-linker-eGFP | This work |
| pMB323 | pRSII406-pif1-m1-I(-5)A-6xHis-6xFlag | This work |
| pMB326 | pRSII406-pif1-4L-6xHis-6xFlag | This work |
| pMB328 | pRSII406-pif1-m1-L9L-6xHis-6xFlag | This work |
| pMB329 | pRSII406-pif1-nuc-6xHis-6xFlag | This work |
| pMB338 | pRSII406-pif1-m2-M107I-M113I-6xHis-6xFlag | This work |
| pMB339 | pRSII406-pif1-m2-M113I-M127I-6xHis-6xFlag | This work |

Supplementary Table S3. **Oligonucleotides for PCR, cloning and sequencing employed in this work.**

| **Alias** | **Name** | **Sequence** |
|---|---|---|
| OMB42 | PIF1 FW (-226) | TTACATTAAGAAAGGCGCGTC |
| OMB43 | PIF1 RV (+70) | GAACTATGTATCTCGGCCTTG |
| OMB55 | delPIF1 FW | TGTAATATTATCCATTGAGCGATTAGCTTACTTGTAT<br>CAATCAATTTTAC CGTACGCTGCAGGTCGAC |
| OMB56 | delPIF1 RV | GATTATTATAGCAGTTTGTATTCTATATAACTATGTG<br>TATTAATATGTACATCGATGAATTCGAGCTCG |
| OMB59 | 5'-TOP3 | CCATTATCTATGGTGGCATGTG |
| OMB60 | delTOP3 FW | CTTCGTCTGTGCGCTCGCTTCAGCTACGTAAAGGT<br>GTATTATAGACAATA CGTACGCTGCAGGTCGAC |
| OMB61 | delTOP3 RV | ATGCAATTAAGCGGAGAGCTTTTTTGAAGACAAAG<br>GGCGGCAAAAACGCC ATCGATGAATTCGAGCTCG |
| OMB67 | PIF1 FW SEQ1 (240) | TGATTGGGAAGAACCTGATTGC |
| OMB68 | PIF1 FW SEQ2 (492) | CAAGAATCCATTAAGACCAGCG |
| OMB69 | PIF1 FW SEQ4 (1021) | TATTGGTGCTTTGGTTGTCG |
| OMB70 | PIF1 FW SEQ3 (1621) | GATCCCTTGGTAAAGTCATCG |
| OMB75 | PIF1 tag FW | TGGTTTCCGACGAACCTCGTGGTCAGGATACCGAA<br>GACCACATCTTAGAATCCGGTTCTGCTGCTAG |
| OMB76 | PIF1 tag RV | GATTATTATAGCAGTTTGTATTCTATATAACTATGTG<br>TATTAATATGTACCCTCGAGGCCAGAAGAC |
| OMB84 | DNA2 FW (-157) | AACTGATCTTACGCTATTTATGGC |
| OMB85 | DNA2 RV (+114) | CGAACCTCAGATGCTAATATTACG |
| OMB87 | delDNA2 FW | GAGTACTCATTTTGTGCAAGCAAACACTGACAATTG<br>AAGAGATCGTCAGGCGTACGCTGCAGGTCGAC |
| OMB88 | delDNA2 RV | TATGCTGTGATAGCTTTCCTGTTATGGAGAAGCTCT<br>TCTTATTCCCCCTGATCGATGAATTCGAGCTCG |
| OMB129 | delNLS-Pif1-FW (623-646) | CATCAAAACTCTGCAGGAAAACG |
| OMB130 | delNLS-Pif1-RV (623-646) | CATGAAGTCAAAAACTGAGGAATCTAGG |
| OMB131 | delNLS-Pif1-FW (650-662) | GCTTCTGATATGAGTACGAGGATGG |
| OMB132 | delNLS-Pif1-RV (650-662) | GTTTTGATGTATGGTCTGCATCAG |
| OMB133 | delNLS-Pif1-FW (814-834) | TCTAATAGTAATCAAGTTCATTCATTGG |
| OMB134 | delNLS-Pif1-RV (814-834) | AGATTGTGTGGTCGCTG |
| OMB208 | delNLS-Pif1-FW (775-803) | CCAGCACCCATATCAGC |
| OMB209 | delNLS-Pif1-RV (775-803) | AAGTTGCTTATAGGCACTTTCG |
| OMB210 | PIF1-775-803-4A-FW | TTAGACTACGCACCAGGC |
| OMB211 | PIF1-775-803-4A-RV | TGCTGCAGCAGCCACTTGCTCATCTGCCTC |

| | | |
|---|---|---|
| OMB212 | PIF1-775-803-5A-FW | CCTGCATATGCAGCTGCTTCCGCATCAGCTTCAAA TTCTCCAGCACCC |
| OMB213 | PIF1-775-803-5A-RV | GCCTGGTGCGTAGTCTAAC |
| OMB228 | AttB1-PIF1-M1-FW | GGGGACAAGTTTGTACAAAAAAGCAGGCTTCGCCC CAAAGTGGATAAGATCAAC |
| OMB238 | PIF1 P.I | TTCTAAGATGTGGTCTTCGGTATCCTGACCACGAG GTTCGTCGGAAACCA TTCGTACGCTGCAGGTCGAC |
| OMB241 | PIF1-M107I-FW | AGGGATAAAAATGTCATGAATTTTATC |
| OMB242 | PIF1-M107I-RV | AATAGGCTTTTTGTCCACC |
| OMB243 | PIF1-M113I-FW | AATTTTATCAATAAAGACAGTCCTTTATCC |
| OMB244 | PIF1-M113I-RV | AATGACATTTTTATCCCTCATAGG |
| OMB245 | PIF1-M127I-FW | TTTAAACCCAGTATAATACAACCAC |
| OMB246 | PIF1-M127I-RV | AATATCGTTCCAGGATAAAGG |
| OMB247 | PIF1-M187I-FW | ATAAATGAAAACGAAAAGAAGAAAATG |
| OMB248 | PIF1-M187I-RV | AATTTCCAAACTTCTCTCTTGAG |
| OMB249 | PIF1-M196I-FW | CAATTTGGAGAAAAGATTGCTG |
| OMB250 | PIF1-M196I-RV | AATTTTCTTCTTTTCGTTTTCATTTATC |
| OMB251 | DIA2 (-287) FW | CATTGATCGTTTGAGTATCTGCG |
| OMB252 | DIA2 (-101) FW | TTGGTAACACTAGCTTTAAACCG |
| OMB253 | DIA2 (+91) FW | AGCTTTCTATATTCTCACCTCC |
| OMB254 | DIA2 (+213) RV | TTGGTAACACTAGCTTTAAACCG |
| OMB261 | PIF1-MET1-Kozak-RV | GTTTAATTGATTGATACAAGTAAGCTAATCG |
| OMB262 | PIF1-MET1-Kozak-FW | ATGCCAAAGTGGATAAGATC |
| OMB263 | PIF1-MET2-Kozak-RV | ACTGAACGATAGTTTGGCGTGC |
| OMB265 | PIF1-MET2-Kozak-FW | ATGTCTAGTCGTGGTTTCAGGTCTAATAAC |
| OMB286 | PIF1-S1 | TTTGTAATATTATCCATTGAGCGATTAGCTTACTTGT ATCAATCAATTTTACATGCGTACGCTGCAGGTCGAC |
| OMB287 | PIF1-S4 | AATGGCCTTCTTGGTATAATATGATTCAATGTTGAT CTTATCCACTTTGGCATCGATGAATTCTCTGTCG |
| OMB288 | PIF1-775-803-9A-RV | GCCTGGTGCGTAGTCTAATGC |
| OMB314 | PIF1-M107I-M113I-RV | AATGACATTTTTATCCCTAATAGGC |
| OMB315 | PIF1-M107I-M113I-FW | AATTTTATCAATAAAGACAGTCCTTTATCC |
| OMB338 | M107-PIF1-pRS-RV | GTAAAATTGATTGATACAAGTAAGC |
| OMB339 | M107-PIF1-FW | ATGAGGGATAAAAATGTCATG |
| OMB340 | NdeI-M107-PIF1-pET28b-FW | ATCGATCATATGAGGGATAAAAATGTCATG |
| OMB341 | M107-PIF1-pET28b-RV | AAGCTTGTCGACTTATTCTAAG |
| OMB342 | M107-PIF1-pAG-RV | GAAGCCTGCTTTTTTGTAC |
| OMB349 | m1045 | CTGCATTTGGCTCCATTTT |
| OMB350 | m1046 | TGTTGTAATGGGGACGGAAA |

| | | |
|---|---|---|
| OMB363 | Telomeric-Probe-Y-FW | GCGTTTGCGTTCCATGACGAG |
| OMB364 | Telomeric-Probe-Y-RV | CTGCCGTGCAACAAACACTAAATC |
| OMB409 | PIF1-N-term P.II | TGTAATATTATCCATTGAGCGATTAGCTTACTTGTATCAATCAATTTTAC CCGCGCGTTGGCCGATTCAT |
| OMB416 | PIF1-WT-I5L-RV | TGTTGATCTTAACCACTTTGGCATGTAAAATTG |
| OMB417 | PIF1-L9L-I12L-IL13L-FW | TTAAATCATTTATTACCAAGAAGGCCATTTATCTG |
| OMB419 | PIF1-L9L-RV | TGTTGATCTTATCCACTTTGG |
| OMB420 | PIF1-L9L-FW | TTAAATCATATTATACCAAGAAGGC |
| OMB421 | PIF1-M1-I5-9-12-13L-RV | TGATCTTAACCACTTTGGGGCCCAAAATTG |
| OMB422 | PIF1-M1-I5-9-12-13L-FW | ACATTAAATCATTTATTACCAAGAAGGCCATTTATCTG |
| OMB427 | PIF1-I(-5)A-RV | ACAAGTAAGCTAATCGCTCAATGG |
| OMB430 | PIF1-M1-I(-5)A-FW | GCTAATCAATTTTGGGCCCCAAAGTGG |

**Supplemental references**


1. Meijer, H. A. & Thomas, A. A. M. Control of eukaryotic protein synthesis by upstream open reading frames in the 5′-untranslated region of an mRNA. *Biochem. J.* **367**, 1–11 (2002).
2. Hernández, G., Osnaya, V. G. & Pérez-Martínez, X. Conservation and Variability of the AUG Initiation Codon Context in Eukaryotes. *Trends Biochem. Sci.* **44**, 1009–1021 (2019).
3. Archer, S. K., Shirokikh, N. E., Beilharz, T. H. & Preiss, T. Dynamics of ribosome scanning and recycling revealed by translation complex profiling. *Nature* **535**, 570–574 (2016).
4. Michel, A. M. *et al.* GWIPS-viz: development of a ribo-seq genome browser. *Nucleic Acids Res.* **42**, D859–D864 (2014).
5. Vega, L. R. *et al.* Sensitivity of yeast strains with long G-tails to levels of telomere-bound telomerase. *PLoS Genet* **3**, e105 (2007).
6. Rothstein, R. J. [12] One-step gene disruption in yeast. in *Methods in Enzymology* vol. 101 202–211 (Academic Press, 1983).

7. Houseley, J. & Tollervey, D. Repeat expansion in the budding yeast ribosomal DNA can occur independently of the canonical homologous recombination machinery. *Nucleic Acids Res.* **39**, 8778–8791 (2011).

8. Wach, A., Brachat, A., Pöhlmann, R. & Philippsen, P. New heterologous modules for classical or PCR-based gene disruptions in Saccharomyces cerevisiae. *Yeast* **10**, 1793–1808 (1994).

9. Longtine, M. S. *et al.* Additional modules for versatile and economical PCR-based gene deletion and modification in Saccharomyces cerevisiae. *Yeast* **14**, 953–961 (1998).

10. Storici, F. & Resnick, M. A. The Delitto Perfetto Approach to In Vivo Site-Directed Mutagenesis and Chromosome Rearrangements with Synthetic Oligonucleotides in Yeast. in *Methods in Enzymology* vol. 409 329–345 (Elsevier, 2006).

11. Alberti, S., Gitler, A. D. & Lindquist, S. A suite of Gateway® cloning vectors for high-throughput genetic analysis in Saccharomyces cerevisiae. *Yeast* **24**, 913–919 (2007).

12. Boulé, J.-B., Vega, L. R. & Zakian, V. A. The yeast Pif1p helicase removes telomerase from telomeric DNA. *Nature* **438**, 57–61 (2005).
